# Antibiotic lethality dictates mycobacterial infection outcomes

**DOI:** 10.1101/2024.10.29.619904

**Authors:** Alexander Jovanovic, Frederick K. Bright, Ahmad Sadeghi, Basil Wicki, Santiago E. Caño Muñiz, Sara Toprak, Loïc Sauteur, Anna Rodoni, Andreas Wüst, Andréanne Lupien, Sonia Borrell, Dorothy M. Grogono, Nicole Wheeler, Philippe Dehio, Johannes Nemeth, Hans Pargger, Rachel Thomson, Scott C. Bell, Sebastien Gagneux, Josephine M. Bryant, Tingying Peng, Andreas Diacon, R. Andres Floto, Michael Abanto, Lucas Boeck

## Abstract

Antibiotic development and treatment focus on bacterial growth inhibition, often with limited success. Here, we introduce Antimicrobial Single-Cell Testing (ASCT), an advanced imaging strategy to assess bacterial killing in real-time. By tracking 140 million bacteria and generating over 20,000 *in vitro* time-kill curves, we can predict *Mycobacterium tuberculosis* treatment outcomes in mice and humans and link strain-specific survival (drug tolerance) in *Mycobacterium abscessus* to clinical responses. Using ASCT, we reveal drug tolerance as a distinct genetically encoded bacterial trait conserved across drugs with similar targets and, via genome-wide associations, uncover molecular mechanisms that govern bacterial killing. This study establishes the technical framework and *in vivo* validation for large-scale bacterial killing assessments to advance our understanding of bacterial survival, antibiotic development and clinical decision-making.

## BACKGROUND

Bacteria have evolved diverse strategies to overcome toxic exposures. The best-known is antibiotic resistance, where genetic modifications alter the target site or change the effective drug concentration at the target, enabling bacterial growth during drug treatment (*1*). Another bacterial strategy involves transient bacterial survival during antibiotic treatment, allowing bacterial regrowth once the drug is cleared. This phenomenon is known as drug tolerance, when the overall population killing is delayed, and persistence when a highly tolerant subpopulation is present (*2*). Despite its decades-long recognition (*3*–*5*), the mechanisms underlying bacterial drug tolerance in human infections, and especially its relevance for antibiotic treatment, remain largely unknown.

Even in the absence of antibiotic resistance, treatment outcomes for bacterial infections – including urinary tract, respiratory and bloodstream infections – are sometimes poor (*6, 7*). This challenge is highlighted in mycobacterial infections, where limited treatment efficacy necessitates prolonged multidrug therapy, often requiring drug combination treatments for several months to years. For drug-susceptible *Mycobacterium tuberculosis*, which caused more than 1 million deaths in 2022 (*8*), it took decades of extensive research and clinical trials to achieve even a modest reduction in treatment duration (from six to four months) (*9*). These extended treatment times are costly, associated with treatment-related toxicity, and increase the risk of non-adherence, potentially driving relapses and transmission. The therapeutic challenge is even more acute in *Mycobacterium abscessus* infections, now one of the most prevalent mycobacterial infections in developed countries (*10*). For *M. abscessus*, no consistently effective drug regimen exists, and treatment outcomes are often poor, with failure rates frequently exceeding 50% despite prolonged treatment with multiple drugs that show *in vitro* activity (*11*).

The poor treatment efficacy against drug-susceptible mycobacteria motivated us to look beyond standard antibiotic susceptibility testing (*12, 13*) to develop an *in vitro* assay that mimics *in vivo* drug responses and infection outcomes. Minimum inhibitory concentrations (MICs) do not address antibiotic killing, which has been challenging to study due to the labour-intensive, low-throughput nature of colony-forming unit (CFU) assays (*14*–*16*). To bridge this gap, we established Antimicrobial Single-Cell Testing (ASCT), a highly scalable method for quantifying antibiotic lethality at the single-cell level. This approach enabled us to examine drug as well as bacterial effects on antibiotic killing, to uncover mechanisms and clinical implications of bacterial survival and death.

## RESULTS

### Antimicrobial Single-Cell Testing captures bacterial killing and survival at massive scale

To quantify bacterial growth and antibiotic killing in bacterial populations and subpopulations at a scale comparable to MIC assessments, we developed Antimicrobial Single-Cell Testing (ASCT; ***Fig. 1***), a workflow that simultaneously tracks the behaviours and the viability of millions of bacterial cells across more than 1,000 different conditions. In ASCT, bacteria are dispensed into multi-well plates, immobilised in agar pads containing propidium iodide (PI) and exposed to drugs (***Supplementary Text, Fig. 1A***). High-content live-cell imaging captures brightfield and fluorescence images of over 10,000 fields every two to four hours for up to seven days, generating up to one million images per experiment. Time-lapses of every field are sequentially analysed using sparse and low-rank decomposition to adjust for spatial and temporal variations in background fluorescence (*17*), supervised random forest classifiers to segment individual bacteria and classify their viability (*18*), drift correction, and single-cell tracking through object position and homology (***Fig. 1B-C, fig S1A-E***) (*17, 18*). To analyse kill kinetics at the population level, single-cell information is pooled and quantified using the area under the time-kill curve as a composite measure of drug tolerance and persistence. While ASCT can also quantify homogenous and heterogenous bacterial growth (e.g., resistance or hetero-resistance), this work focuses on the determinants and consequences of antibiotic killing.

**Figure 1.**
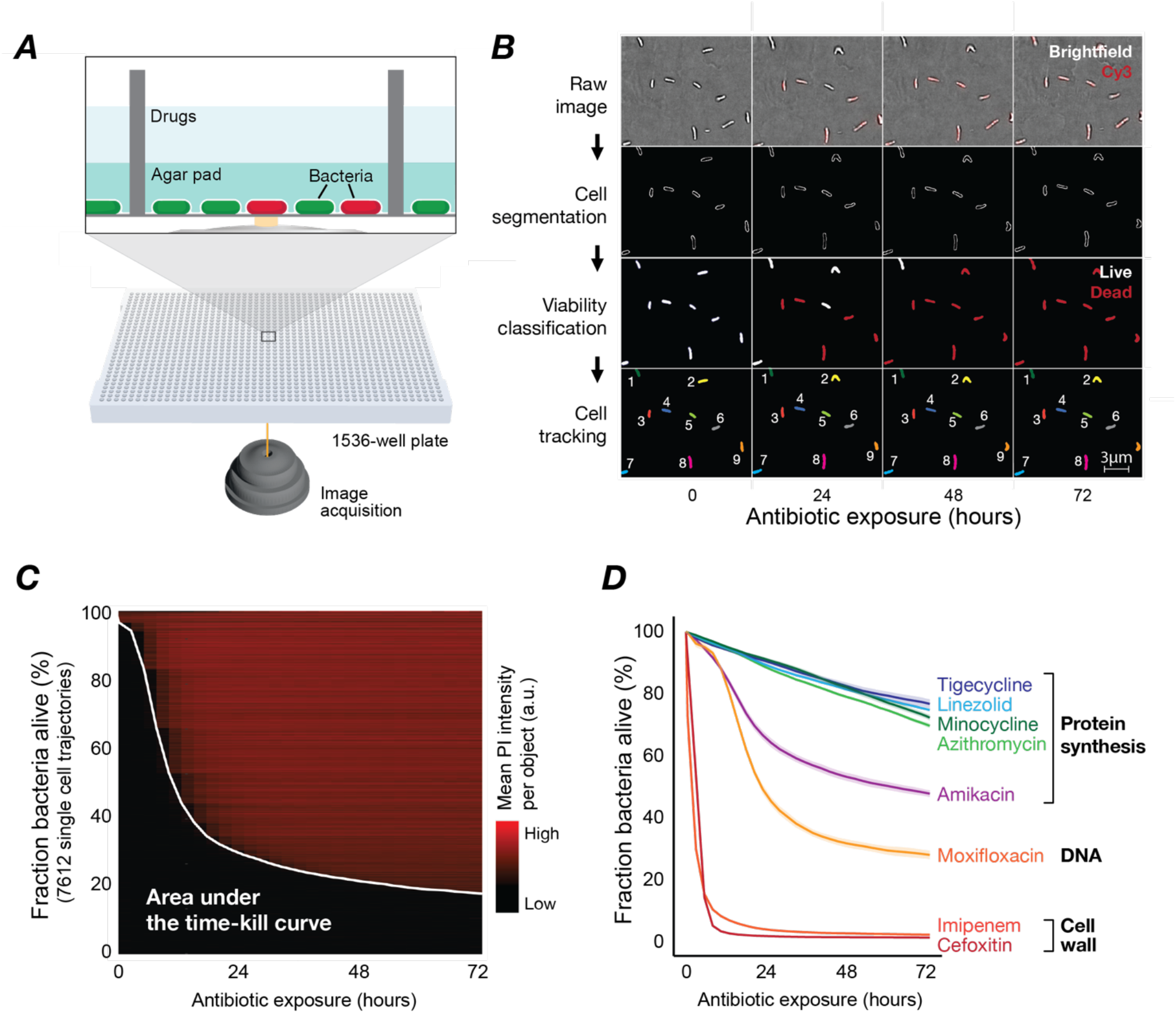
Antimicrobial Single-Cell Testing enables high-throughput assessments of bacterial trajectories. **(A)** Illustration of the ASCT workflow. Bacteria (dead cells in red) are immobilised within agar pads (in wells of a 1536-well plate) and exposed to drugs. Live-cell imaging (brightfield and fluorescence) of nine fields per well and up to 1536 wells is performed in 2-to 4-hour intervals from the plate bottom. **(B)** Schematic of the automated image analysis. Raw brightfield images (cell morphology) and fluorescence images (quantifying propidium iodide [PI] accumulation via Cy3 settings) are processed to segment cells (lines indicate cell borders), classify bacterial viability (white: viable; red: dead) and track individual cells (track IDs indicated by numbers). **(C)** Example of 7,612 *M. abscessus* single-cell trajectories (from a single imaging well) upon tigecycline exposure with PI intensity over time. Bacterial trajectories are ordered by the time of PI conversion. The white line represents the proportion of live bacteria across 29 time points and reveals population time-kill kinetics. The area under the time-kill curve (AUC) was used as a measure of overall killing. **(D)** ASCT-derived time-kill curves (mean ± SEM of three replicates) of a clinical *M. abscessus* isolate exposed to multiple antibiotics, with corresponding targets indicated.

The viability of each bacterium was assessed using PI, a cell-impermeable fluorescent dye. PI accumulation within bacteria, which only occurs in the presence of membrane disruption, was used as a proxy marker for bacterial death, as previously demonstrated in various bacterial species (*19*– *21*). To validate this method, we treated *M. abscessus* with a number of individual antibiotics for 24 hours and tracked more than 30,000 single bacteria for a further 24 hours post-antibiotic washout. Only one PI-positive bacterium resumed growth (likely a misclassified duplet of a live and dead bacterium), while 1-11% of the PI-negative bacteria regrew in antibiotic-free media (***fig. S1F***), confirming that PI accumulation is a reliable marker of bacterial death. ASCT-based time-kill kinetics in *M. abscessus* demonstrated reproducible and drug-specific patterns that aligned with bacteriostatic and bactericidal drug classes and remained relatively stable when using different gel volumes or gel concentrations (***Fig. 1D, fig. S1G-H***).

### Time-kill kinetics of *M. tuberculosis* drug combinations predict *in vivo* outcomes

We first investigated whether *in vitro* drug effects on bacterial killing relate to *in vivo* outcomes observed in mice or humans. Specifically, we analysed the potential of ASCT to identify more effective treatments for tuberculosis. We tested the two avirulent *M. tuberculosis* strains H37Ra and mc^2^7000, during exponential growth in nutrient-rich and under starvation conditions across 65 drug combinations. Each drug was dosed at the maximum blood concentration (C_max_) achievable during therapeutic dosing in humans. We revealed that only drug regimens containing isoniazid, rifampicin or ethambutol (all drugs included in the standard *M. tuberculosis* regimen) were favourable in killing exponentially-growing *M. tuberculosis* (***fig. S2A***,***C-D***). In contrast, regimens including bedaquiline, clofazimine or pretomanid outperformed other drugs or combinations under starvation conditions (***Fig. 2A-B, fig. S2B***).

**Figure 2.**
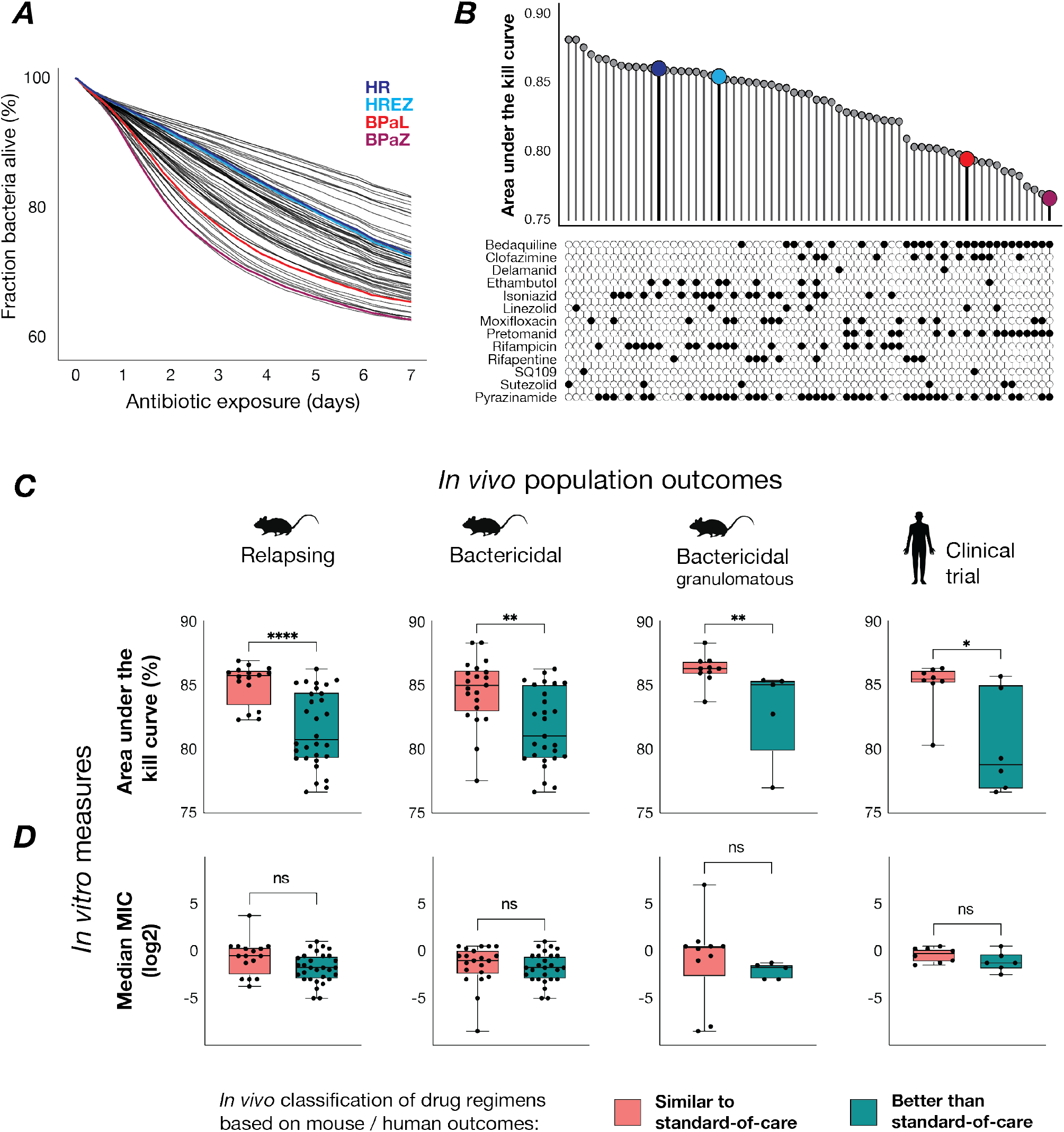
Time-kill kinetics predict *in vivo* outcomes of *M. tuberculosis* drug regimens. **(A)** ASCT-based time-kill kinetics of starved *M. tuberculosis* (PBS-starved mc^2^7000) exposed to 65 drug regimens, with the following regimens highlighted: isoniazid-rifampicin (HR; dark blue), isoniazid-rifampicin-ethambutol-pyrazinamide (HREZ; light blue), bedaquiline-pretomanid-linezolid (BPaL; red), and bedaquiline-pretomanid-pyrazinamide (BPaZ; purple). **(B)** Areas under the kill curve for *M. tuberculosis* drug regimens averaged across three *M. tuberculosis* starvation conditions. **(C-D)** Drug regimens were previously classified as similar or better than standard of care (SOC) based on the performance in relapsing mouse models (RMM), bactericidal mouse models (BMM) of common mouse strains, the granulomatous C3HeB/FeJ mouse strain and clinical studies (*22, 23*). **(C)** Time-kill curves (AUC averaged across three *M. tuberculosis* starvation models) of “similar to SOC” drug regimens were compared with “better than SOC” regimens using the Mann-Whitney U test. Each dot represents a drug regimen. Boxplots show the median, interquartile range and total range of AUC values. **(D)** Median minimum inhibitory concentrations (MICs) of drug regimens, calculated from the MICs of individual drugs (averaged from mc^2^7000 and H37Ra strains), were compared between “similar to SOC” and “better than SOC” regimens. (RMM: n = 46, BMM common strains: n = 48, BMM C3HeB/FeJ: n = 15, Clinical bactericidal activity: n = 14). * P < 0.05, ** P < 0.01, **** P < 0.0001, ns: non-significant.

To benchmark ASCT, we used previously established classifications of the *in vivo* efficacy of *M. tuberculosis* drug regimens, categorised as either similar or superior to standard of care (SOC), based on a review of mouse and clinical studies (*22, 23*). While time-kill kinetics of exponentially growing *M. tuberculosis* did not discriminate between drug combinations, we found strong associations between *in vivo* outcomes and *in vitro* killing under starvation conditions (***Fig. 2C, fig. S3A***). These findings suggest that bacteria with reduced metabolic activity, commonly found in intra- and extracellular infection niches, are more challenging to clear and likely responsible for long-term infection outcomes (*24*–*28*). For example, killing of pantothenate-starved mc^2^7000 (the auxotrophic strain H37Rv ΔpanCD ΔRD1) was highly associated with outcomes in the relapsing mouse model, the bactericidal model of common and the C3HeB/FeJ mouse strains, mirroring different outcomes (bacteriologic burden and relapses) and host pathobiology (*29, 30*). Moreover, mc^2^7000 killing upon pantothenate starvation was associated with clinical outcomes (bactericidal activity) in phase 2 clinical studies. Similar results were obtained after 14-day phosphate-buffered saline (PBS) starvation of the mc^2^7000 and H37Ra *M. tuberculosis* strains. Overall, 11 out of 12 starvation conditions reached statistical significance. In contrast, the association of the lowest MIC, median MIC, or mean MIC (of the two most potent drugs) within each combination with *in vivo* outcome measures in mice or humans was poor (***Fig. 2D, fig. S4***). The performance of ASCT for predicting outcomes of clinical studies using *M. tuberculosis* starvation models yielded an area under the receiver operating curve between 0.76 and 0.94 (***fig. S3B***), suggesting that the killing activity of *M. tuberculosis* regimens upon starvation is a critical *in vitro* marker which translates into *in vivo* outcomes, and highlighting the potential of ASCT for prioritising antibiotic combination treatments.

### Drug tolerance is a distinct measure of antibiotic efficacy

We next explored the bacterial factors that influence bacterial killing in *M. abscessus*. We obtained a single bacterial isolate from 405 patients, predominantly individuals with Cystic Fibrosis, with *M. abscessus* lung infection across Europe and Australia (***Fig. 3A***). This allowed us to evaluate whether phenotypic and genetic bacterial features within a naturally evolved bacterial species affect antibiotic killing profiles (i.e., drug tolerance) (*31, 32*). Using ASCT, we generated time-kill profiles for each clinical isolate and the *M. abscessus* laboratory strain against eight drugs at two concentrations, each in triplicate. In total, we assessed 18,244 time-kill curves and tracked approximately 130 million bacterial cells over 29 time points and 72 hours. Isolate-drug pairs that exhibited bacterial growth (i.e., likely reflecting drug resistance) or did not meet quality criteria were excluded from analyses (***Supplementary Text***). Our findings revealed highly diverse but reproducible time-kill kinetics, indicating that specific strain characteristics modulate antibiotic killing within *M. abscessus* (***Fig. 3B***).

**Figure 3.**
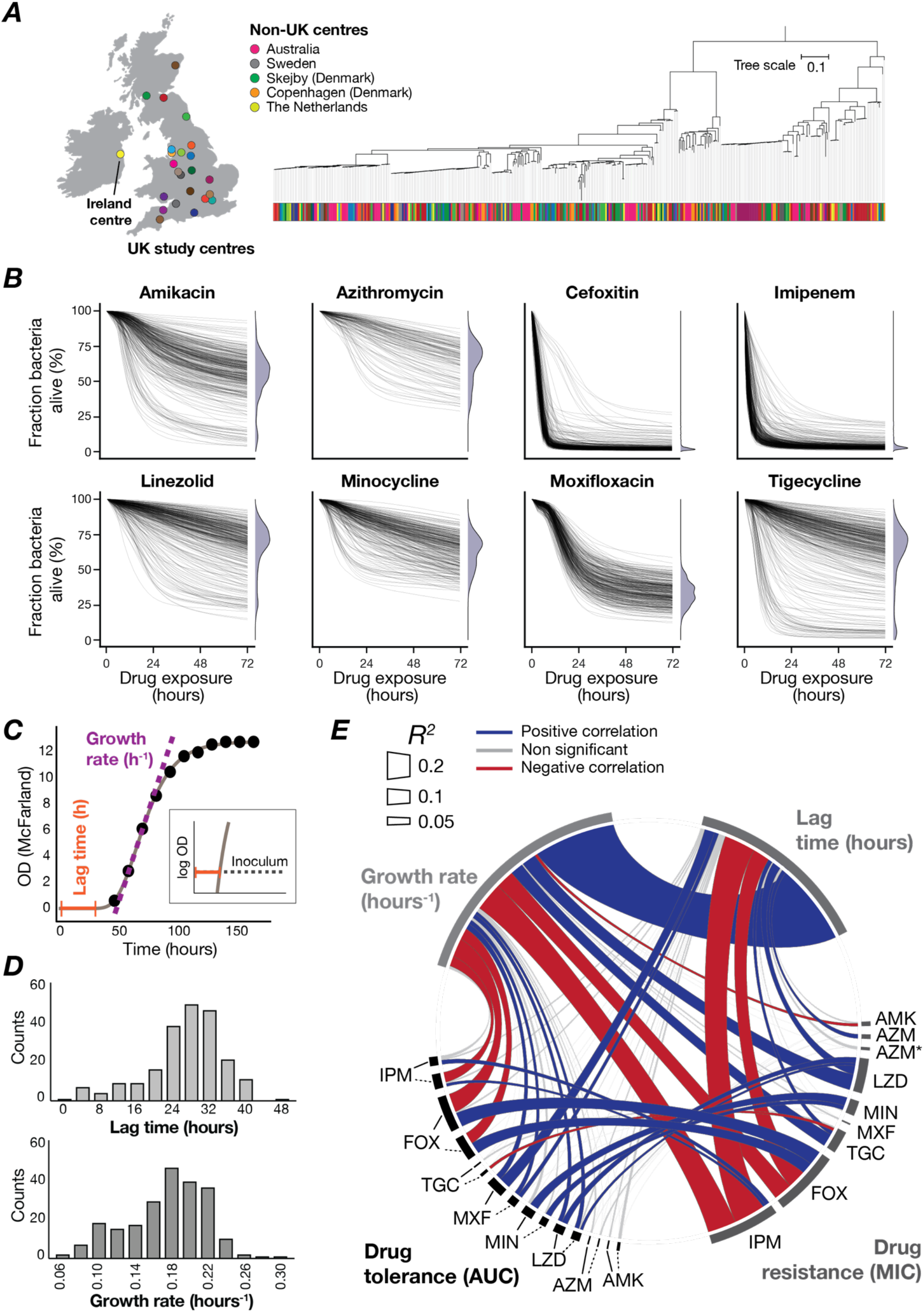
Antibiotic tolerance is a distinct measure of antibiotic efficacy. **(A)** Study centres in the United Kingdom, Ireland, mainland Europe and Australia. The maximum likelihood phylogenetic tree of 376 *M. abscessus* isolates obtained from different patients, with each centre indicated. **(B)** Time-kill curves of 406 *M. abscessus* isolates across eight antibiotics (the lower drug concentrations are shown), with viability distributions at 72 hours, excluding growing isolates. **(C)** Gompertz function fitting to optical density (OD) measurements (black dots), illustrating the determination of bacterial growth rates and lag times. The insert highlights the estimation of lag time in logarithmic scale. **(D)** Distributions of lag times and growth rates from 226 fitted *M. abscessus* growth curves. **(E)** Pearson correlation analyses between drug tolerance (high drug concentration: continuous line, low concentration: dashed line), bacterial growth rate, lag time, and the corresponding minimum inhibitory concentration (MIC). The line colour indicates the direction of the correlation, the line width represents the corresponding R^2^ value. AUC indicates the area under the kill curve, AMK amikacin, AZM azithromycin, AZM* inducible azithromycin resistance assessed on day 14, FOX cefoxitin, IPM imipenem, LZD linezolid, MIN minocycline, MXF moxifloxacin, TGC tigecycline.

Drug tolerance has been linked to slow bacterial growth or prolonged lag times, associated with metabolic dormancy (*33, 34*). To evaluate how bacterial replication affects antibiotic killing, we measured optical densities of 226 *M. abscessus* strains during planktonic growth in nutrient-rich conditions and fitted Gompertz functions to estimate each isolate’s lag time and growth rate (***Fig. 3C-D***). Overall, we identified only weak associations between drug tolerance and bacterial replication. Apart from revealing a strong direct correlation between growth rate and lag time, likely reflecting a trade-off between bacterial growth and adaptation (***Fig. 3E, fig. S5*** (*35*)), we identified correlations between growth rates or lag times and drug tolerance phenotypes. For instance, drug tolerance to imipenem and cefoxitin, which target cell wall synthesis specifically during bacterial replication, correlated with slow growth but not lag time. In contrast, tolerance to moxifloxacin (targeting DNA) was associated with longer lag times only. These associations were weak, explaining only up to 13% of the variation in the drug tolerance phenotype, suggesting that replication patterns of these bacterial populations do not drive drug tolerance. Similar correlations (R^2^ up to 0.2) were observed between growth rates or lag times and drug resistance. We also found weak associations (R^2^ up to 0.11) between drug tolerance and resistance phenotypes (especially for cefoxitin and minocycline), mainly driven by outliers with MICs closer to the drug concentration used in ASCT. Using a non-parametric test (Spearman correlation), most of these associations disappeared (***fig. S5***), supporting that drug tolerance is a marker of antibiotic efficacy independent of drug resistance (*36*).

### Drug tolerance is heritable

While drug resistance is widely recognised as a genetic trait, drug tolerance has traditionally been viewed as a phenotypic, mostly non-genetic, characteristic (*37*). We employed whole-genome sequencing and estimated the proportion of phenotypic variance explained by genetic variability, known as heritability (*38*). We mapped 1.3 million *M. abscessus* unitigs (sequences of variable length), which capture most genomic variation, such as single nucleotide polymorphisms, insertions, deletions and gene presence-absence, to respective phenotypes using linear mixed models (*39, 40*). Drug resistance phenotypes, particularly macrolide MICs, were primarily driven by bacterial genetics, while resistance phenotypes of unstable drugs or drugs with less variable MIC were less heritable (***Fig. 4A***). All tolerance phenotypes demonstrated substantial heritability (32 to 97%), which was different from random chance (Student’s t-test P < 2.2x10^-16^; mean heritability of random chance: 1.1%), indicating that most variability in time-kill kinetics is explained by genetic variance. Mapping these phenotypes to the *M. abscessus* phylogeny (***Fig. 4B, fig. S6***), we observed that some of the high- or low-tolerance phenotypes evolved recently in parallel, resulting in homoplastic traits, while others were shared between phylogenetically related isolates, indicating inherited clades. One such example is a tigecycline low-tolerance clade within the dominant circulating clone of *M. abscessus massiliense* (*41*). Many of these isolates harbour high-level mutational aminoglycoside and macrolide resistance and have been associated with increased virulence (*32*). Our finding of tigecycline low-tolerance highlights a potential vulnerability in these extremely challenging-to-treat *M. abscessus* isolates.

**Figure 4.**
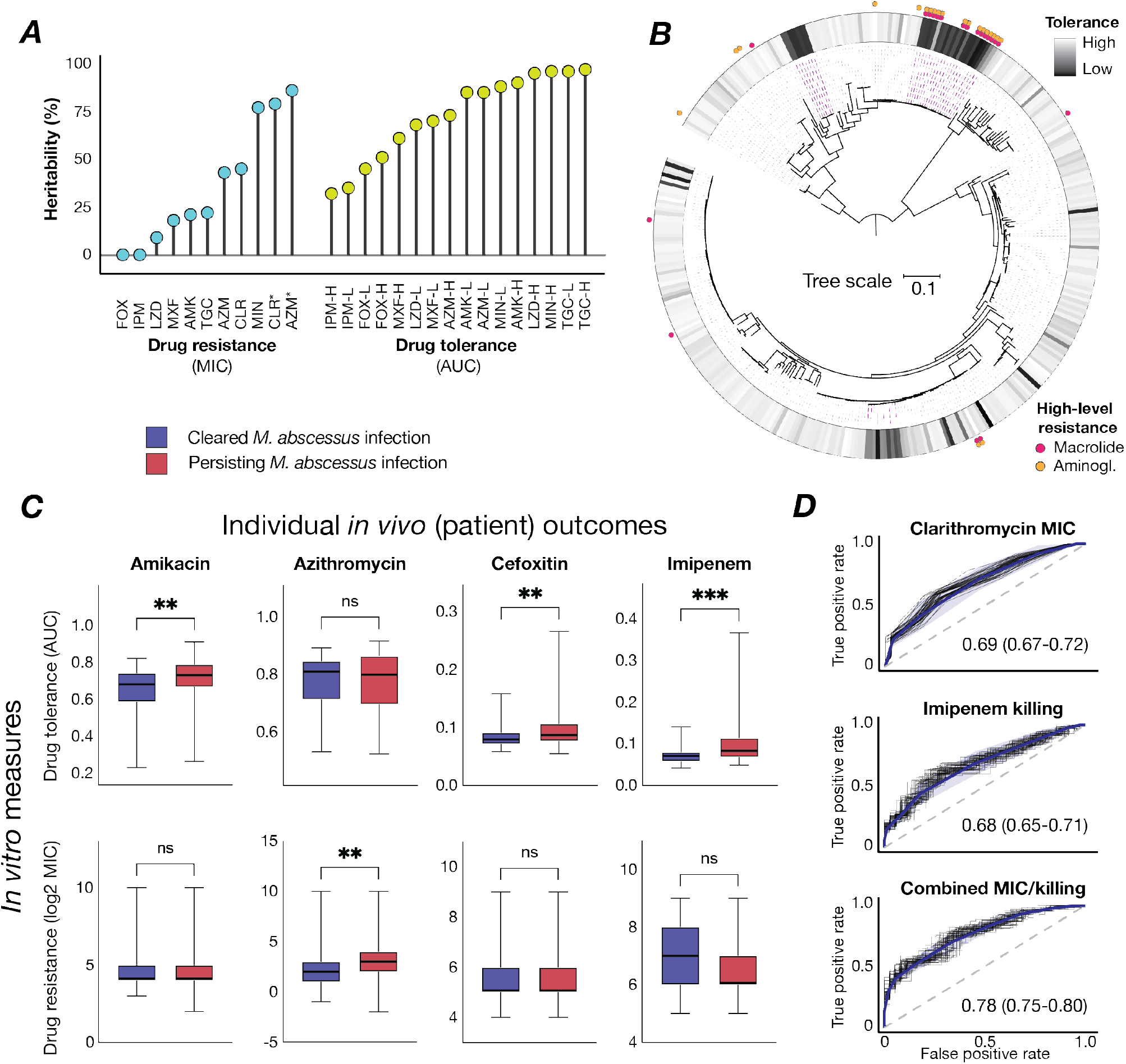
Drug tolerance is driven by the genetic background and correlates with clinical outcomes. **(A)** Heritability estimates for drug resistance (MIC) and drug tolerance (AUC) phenotypes based on 1.3 million genetic *M. abscessus* variants (unitigs) derived from whole-genome sequencing. **(B)** Maximum-likelihood phylogenetic tree of 353 *M. abscessus* isolates, aligned with the tigecycline tolerance heatmap (high concentration) and mutational macrolide and aminoglycoside resistance. Purple lines highlight clades with low tigecycline tolerance. **(C)** Comparison of drug tolerance (AUC, low concentration) and drug resistance (MICs) between *M. abscessus* isolates from patients with poor versus favourable clinical outcomes, using the Mann-Whitney U test. Boxplots show the median, interquartile range and total range of AUC values. **(D)** Logistic regression analysis predicting clinical outcomes based on drug resistance (clarithromycin MIC) and drug tolerance (imipenem at low concentration), individually and in combination. Fifty receiver operating characteristic (ROC) curves were generated by randomly selecting 80% of the samples for each condition. The blue line represents the mean ROC curve, and the shaded area one standard deviation. The mean area under the ROC curve (AUC-ROC) and 95% confidence intervals are shown. AMK indicates amikacin, AZM azithromycin, AZM* inducible azithromycin resistance, CLR clarithromycin, CLR* inducible clarithromycin resistance, FOX cefoxitin, IPM imipenem, LZD linezolid, MIN minocycline, MXF moxifloxacin, TGC tigecycline, L refers to low drug concentration, and H to high drug concentration.

### Drug tolerance predicts infection outcomes

We then investigated whether genotype-specific kill kinetics influence individual patient outcomes, given the limited direct evidence linking drug tolerance to treatment failures (*42*). Our analysis revealed that, while only macrolide MICs were linked to clinical outcomes (***fig. S7A-B***), amikacin, cefoxitin and imipenem tolerance (assessed at both low and high drug concentrations) correlated with *M. abscessus* clearance in patients (***Fig. 4C, fig. S8***). Specifically, rapid killing was associated with favourable clinical outcomes, whereas high tolerance was associated with poor clinical outcomes. Adding macrolide resistance to drug tolerance enhanced the prediction of clinical outcomes from an AUC-ROC of 0.68 to 0.78 (DeLong’s test P = 0.034, ***Fig. 4D***), underscoring that drug tolerance is an independent and relevant marker of antibiotic efficacy.

### Drug tolerance is target specific

By applying principal component analysis to 5,056 *M. abscessus* time-kill curves and mapping antibiotics in drug tolerance space, we identified drug clusters related to the drug’s mode of action (***Fig. 5A***). These clusters indicate that bacterial survival mechanisms are linked to the drug’s target, likely driven by shared downstream effects, such as conserved stress responses that modulate antibiotic killing (*43*). To explore the genetic basis of drug tolerance, we mapped approximately 280,000 *M. abscessus* genotypes (single nucleotide polymorphisms, insertions and deletions) extracted from whole-genome sequences of each isolate to drug tolerance phenotypes, using mixed effect models corrected for population structure. This phenogenomic analysis identified multiple genotypes highly associated with drug tolerance phenotypes (***Fig. 5B***). The top identified genes were commonly shared within groups of antibiotics targeting protein synthesis or the cell wall. We further examined *MAB_0233*, a putative phage tail tape measure protein, in which several deletions were associated with drug tolerance to drugs targeting protein synthesis (***Fig. 5C***). As such, *MAB_0233* could facilitate as a membrane-associated protein intracellular drug accumulation or enhance cellular toxicity via drug-induced mistranslation. Using oligonucleotide-mediated recombineering followed by Bxb1 integrase targeting (ORBIT (*44*)), we generated a *MAB_0233* knockout strain in *M. abscessus*, which showed increased tolerance for antibiotics targeting translation (amikacin, tigecycline and linezolid), but not the cell wall, while MICs remained stable (***Fig. 5D, fig. S9A-B***). This effect on killing was lost following *MAB_0233* complementation.

**Figure 5.**
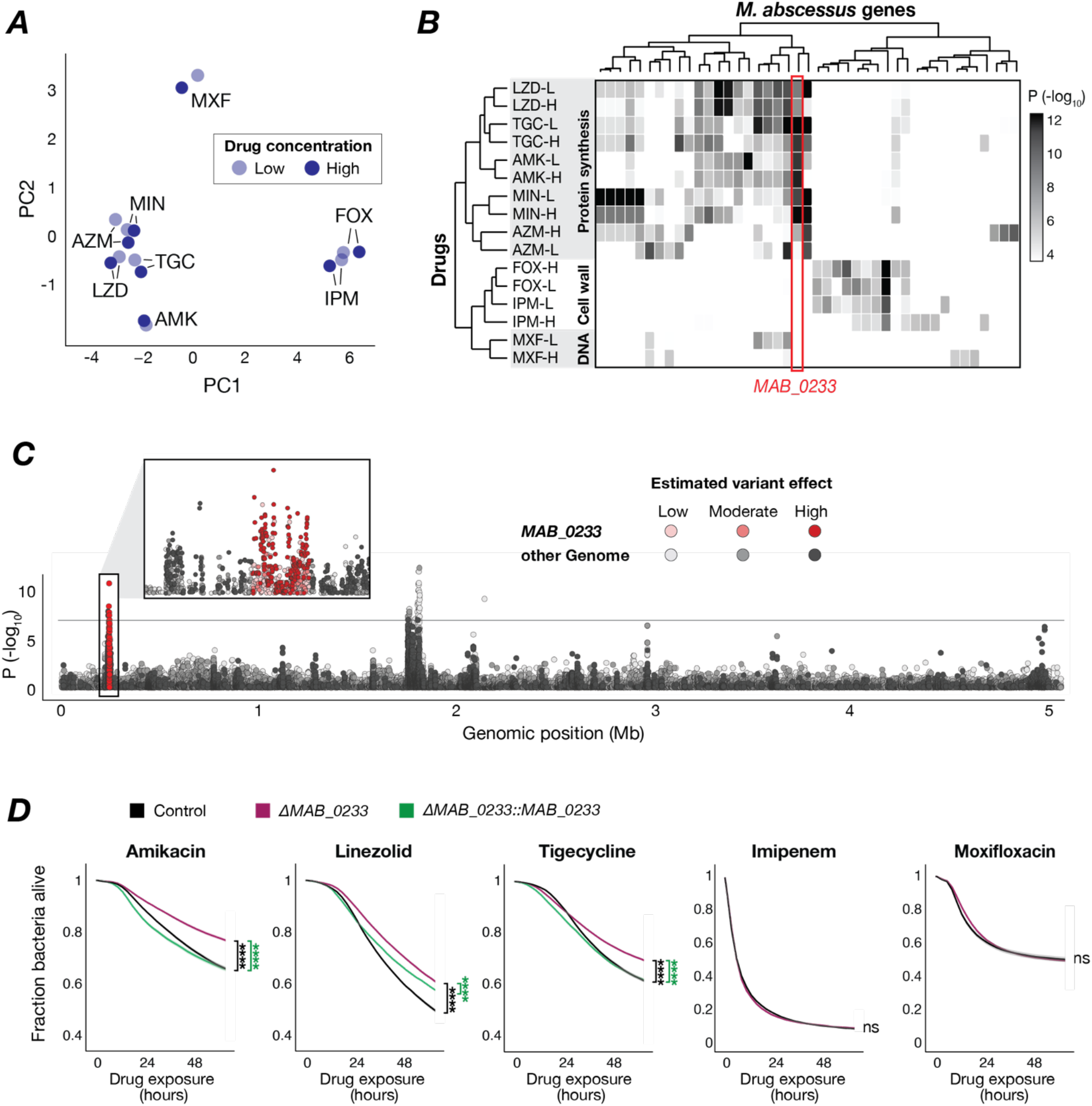
Drug tolerance is target specific. **(A)** Principal component analysis (PCA) of eight drugs at two concentrations in drug tolerance space, using a Spearman correlation matrix of drug tolerance measures from 350 clinical *M. abscessus* isolates. **(B)** Genome-wide association study (GWAS) results for 16 drug tolerance phenotypes, highlighting the five most associated genes (i.e., the top moderate or high effect genotypes) per phenotype. The heatmap shows the association of these 43 genes (top genotype per gene) with tolerance phenotypes using a mixed-effects model corrected for population structure (two-sided Wald test). Associations with P > 0.0001 are white. **(C)** Manhattan plot of 272,351 *M. abscessus* genotypes and their association with amikacin (at low concentration) kill kinetics (two-sided Wald test), with the Bonferroni correction threshold for multiple hypothesis testing (black line, 1.7*10^-7^). Multiple variants in *MAB_0233* were strongly associated with bacterial killing during amikacin exposure (insert). **(D)** Time-kill kinetics of control, *MAB_0233* knockout mutant and the complemented strain, shown as mean ± SEM. AUC values of Δ*MAB_0233 were* compared with the control strain or Δ*MAB_0233::MAB_0233* using the Mann-Whitney U test. AMK indicates amikacin, AZM azithromycin, FOX cefoxitin, IPM imipenem, LZD linezolid, MIN minocycline, MXF moxifloxacin, TGC tigecycline. L refers to low drug concentration, and H to high drug concentration. **** P < 0.0001, ns: non-significant.

## DISCUSSION

Antibiotics that demonstrate efficacy against bacteria *in vitro* commonly fail to achieve clinical success, highlighting a critical disconnect between microbiological assessments and patient outcomes. This gap underscores a major challenge in antibiotic development and patient care. We developed Antimicrobial Single-Cell Testing, a strategy that massively scales up bacterial viability assessments to address a previously underexplored characteristic of antibiotic activity and reveal that insufficient bacterial killing - rather than just resistance - is likely a key factor and, therefore, an *in vitro* predictor of drug trial failures and inconsistent clinical outcomes.

We applied ASCT to investigate the time-kill kinetics of multiple *M. tuberculosis* drug regimens across different strains and experimental conditions. Our findings revealed that, while combinations including the standard drugs isoniazid, rifampicin, and ethambutol were most lethal under exponential growth conditions, the killing efficacy under bacterial starvation predicted *in vivo* outcomes. Antibiotic lethality in these starvation conditions, likely reflective of metabolic inactivity in intra- and extracellular infection niches (*45*), correlated strongly with outcomes in multiple *in vivo* models, including the relapsing and bactericidal mouse models and phase 2 clinical trials. Regimens containing the newer drugs bedaquiline and pretomanid, or the repurposed clofazimine, demonstrated superior killing of starved *M. tuberculosis* and outperformed other combinations *in vivo*. Given the high attrition rates in antibiotic development and clinical trials (*46, 47*), there is an urgent need for more efficient strategies to prioritise therapeutic regimens, which is an increasingly challenging task, considering the growing number of active compounds and possible combinations (*48*). By enabling the evaluation of thousands of drug combinations *in vitro* – with demonstrated relevance *in vivo* – ASCT has the potential to identify more effective and much shorter treatment regimens for *M. tuberculosis*. This advancement could accelerate global health initiatives, such as the World Health Organisation’s goal to eliminate most tuberculosis burden by 2035 (*8, 49*).

Highly effective antibiotics are the cornerstone of successful antibiotic treatments, yet clinical outcomes vary widely, with some patients responding to short courses while others fail despite prolonged multidrug treatment (*50, 51*). We show that antibiotic killing is highly variable across clinical *M. abscessus* isolates, challenging the prevailing view that bacterial killing is solely determined by the drugs bactericidal properties. This variability in killing, different from drug resistance, suggests drug tolerance as a fundamental bacterial trait that may greatly affect treatment efficacy. Notably, there exists scarce evidence that drug tolerance, as opposed to limited drug penetration or reinfections, is underlying antibiotic treatment failures (*52*–*54*). We found that *M. abscessus* strains poorly killed by amikacin, cefoxitin or imipenem (in high and low drug concentrations) were linked to worse clinical outcomes in individual patients, providing some of the most robust evidence to date that drug tolerance is a critical determinant of treatment success. Presumably, due to limited overall killing or infrequent clinical use, such associations were not present in other drugs. Moreover, we show that drug tolerance – like resistance – is largely determined by the bacterial genetic background and, thus, a heritable and evolvable bacterial phenotype. For example, within the subspecies *M. abscessus massiliense*, we identified a low-tolerance clade among highly resistant strains, revealing vulnerabilities for enhancing bacterial clearance.

We used PI as a crude single-cell readout of bacterial viability and confirmed that PI-positive cells do not regrow and thus can be considered dead (*21*). However, the exact viability state of PI-negative cells, which do not regrow after antibiotic washout, remains less clear. Tolerance phenotypes were not explained by slow bacterial growth or extended lag times but instead clustered with the drug’s mode of action, implying shared survival pathways of drugs targeting protein synthesis or the cell wall. Using phenogenomic analysis, we identified numerous genes and genotypes likely conferring high- or low-tolerance phenotypes and validated via gene knockout and complementation that a phage protein modulates bacterial killing. These tolerance mechanisms represent potential targets for sterilising antibiotic treatments, while tolerance genotypes could be integrated into diagnostics, analogous to molecular susceptibility testing. Presumably, molecular tolerance testing, or tolerance phenotyping via ASCT, could facilitate individualised, tolerance-tailored antibiotic regimens and help improve patient outcomes beyond the development of new drugs and regimens.

Our study focused on the killing of bacterial populations, but the capabilities of ASCT extend far beyond this application. By simultaneously tracking millions of single bacteria across hundreds of conditions, ASCT enables the high-throughput screening of bacterial growth and killing, the analysis of population heterogeneity, and the study of single-cell fates. While validated in mycobacteria, this approach is likely applicable to other bacterial species and multidrug-resistant pathogens, offering broad utility in microbiology. ASCT could transform our understanding of bacterial survival and death, bridge the gap between *in vitro* measures and *in vivo* realities, and thereby advance drug development and personalised infection treatments.

## ACKNOWLEDGEMENTS

We want to thank Bettina Schulthess and Peter Sander for training in MIC assessments; Andrej Trauner, Daniel Pinschewer, Mattia Zampieri, Alexander Harms, Damien Portevin, Bree Aldridge and Urs Jenal for expert input; Rubén M. Cabezón, Iñaki M. de Ilarduya, Guillermo Losilla and Martin Jacquot for computational support; Julie Sollier for critical reading and editing of the manuscript; the DBM imaging core facility; and Michael Roth and Michael Tamm for lab space. Calculations were performed at sciCORE scientific computing center at the University of Basel. L.B. received funding from the Swiss National Science Foundation (grant no. 177799, 185792, 215557), Bangerter-Rhyner Foundation, Goldschmidt Jacobson Foundation, Helmut Horten Foundation, Swiss Society for Pneumology and the Cloëtta Foundation. L.B. was supported as part of the NCCR AntiResist, a National Center of Competence in Research, funded by the Swiss National Science Foundation (grant no. 180541).

## AUTHOR CONTRIBUTIONS

Conceptualisation: L.B.; Data curation: A.J., L.B.; Formal analysis: A.S., A.J., L.B.; Funding acquisition: L.B.; Investigation: A.S., A.J., F.K.B., B.W., S.E.C.M, S.T., L.S., A.R., A.W., A.L., L.B.; Methodology: A.S., A.J, F.K.B., L.B.; Project administration: L.B.; Software: A.S., A.J., T.P., L.B.; Resources: D.M.G., N.W., P.D., J.N., H.P., R.T., S.C.B., S.G., J.M.B., A.D., R.A.F., M.A., L.B.; Supervision: L.B.; Validation: A.S., A.J., F.K.B., L.B.; Visualisation: A.S., A.J., S.E.C.M, L.B.; Writing - original draft: L.B.; Writing - review and editing: A.S., A.J., F.K.B., B.W., S.E.C.M, S.T., L.S., A.R., A.W., A.L., D.M.G., N.W., P.D., J.N., H.P., R.T., S.C.B., S.G., J.M.B., T.P., A.D., R.A.F., M.A., L.B.

