## Supplement for "Antibiotic lethality dictates mycobacterial infection outcomes"

**SUPPLEMENTARY TEXT**

#### Sample collection

We used the *Mycobacterium abscessus* laboratory strain ATCC-19977, the avirulent *Mycobacterium tuberculosis* strain H37Ra and the auxotrophic *M. tuberculosis* strain mc$^2$7000 (H37Rv ΔpanCD ΔRD1) (*1*). Clinical *M. abscessus* isolates were obtained from respiratory samples (sputum or bronchoalveolar lavage fluid) collected from patients with pulmonary *M. abscessus* infection as described previously (*2*). Included patients came from all major Cystic Fibrosis centres in the United Kingdom, the Republic of Ireland (Dublin), Sweden (Gothenburg), Denmark (Copenhagen and Skejby), the Netherlands (Nijmegen) and Australia (Queensland). *M. abscessus* isolates were retrieved from the original mycobacterial growth indicator tubes (MGIT) or, if otherwise unavailable, from sub-cultured isolates. The study was approved in England and Wales by the National Research Ethics Service (12/EE/0158) and the National Information Governance Board (ECC 3-03 (f)/2012) and in other centres by respective local review boards.

#### Bacterial growth assessments

Planktonic growth rates and the lag times of *M. abscessus* isolates were assessed following suspension in nutrient-rich media. Approximately $10^5$ CFUs *M. abscessus* were inoculated in 2.5 ml of Middlebrook 7H9 (supplemented with 0.4 % glycerol, 10 % OADC [oleic acid, albumin, dextrose, catalase] and 0.05 % Tween 80) in 15 ml glass tubes, which were then incubated at 37°C with 150 rpm orbital shaking. Optical densities ($OD_{565}$) were measured with a densitometer (DEN-1B; Biosan) and quantified in McFarland units. $OD_{565}$ readings were performed every 12 hours for a minimum of five days until the readings stabilised for at least 24 hours (change in McFarland less than 0.5). Background corrected $OD_{565}$ measurements were used to fit Gompertz functions to estimate growth rates and lag times, where N is the number of bacteria at a given time (t), with A representing carrying capacity (maximum population size), B the initial growth factor affecting displacement along the y-axis and C, the growth rate constant (*3*).

$$N(t) = \; Ae^{-Be^{-Ct}}$$

Given that conventional formulas are imprecise when considering that the growth rate is a proportion of the carrying capacity per time (*4*), we calculated the growth rate (μ) with the following equation:

$$\mu = C \; \frac{lambertW(\frac{A}{eN_0} ln\left(ln\left(\frac{A}{N_0}\right)\right))}{ln\left(ln\left(\frac{A}{N_0}\right)\right)}$$

The lag time (λ) was calculated with:

$$\lambda = -\,ln\left(ln\left(\frac{A}{N_0}\right)/B\right)/C$$

#### Drug susceptibility testing

Antibiotic resistance was quantified with minimum inhibitory concentrations (MICs), following the Clinical Laboratory Standards Institute (CLSI; M24 3rd edition) guidelines (*5*, *6*). B.W. received training at the National Center for Mycobacteria, Zürich (Switzerland), to align with standard clinical microbiology practices. *M. abscessus* isolates were cultured in Middlebrook 7H9, supplemented with 0.4 % glycerol, 10 % OADC and 0.05 % Tween 80, for three days and then diluted in phosphate-buffered saline (PBS) to a McFarland standard of 0.5. A 140 µl aliquot of this suspension was mixed into 14 ml cation-adjusted Mueller-Hinton broth (CAMHB) to achieve an approximate density of $10^6$ CFUs / ml. Antibiotics were tested in log2-fold concentrations steps at the following ranges: amikacin (0.25 – 512 µg/ml), azithromycin (3.9 ng/ml – 512 µg/ml), cefoxitin (0.5 – 256 µg/ml), clarithromycin (3.9 ng/ml – 512 µg/ml), clofazimine (7.8 ng/ml – 16 µg/ml), imipenem (0.13 – 256 µg/ml), linezolid (0.25 – 128 µg/ml), minocycline (15.6 ng/ml – 256 µg/ml), moxifloxacin (15.6 ng/ml – 32 µg/ml), and tigecycline (15.6 ng/ml – 8 µg/ml). For each experiment CAMHB was freshly prepared. Antibiotics and CAMHB were added to 96-well plates to reach a volume of 50 µl per well. After mixing, the 50 µl bacterial suspension was added to each well to obtain a final bacterial concentration of $\sim 5*10^5$ CFUs/ml. Each plate included growth and media control wells. *Mycobacterium peregrinum* (ATCC-700686) MICs were assessed as quality control in each experimental batch. Plates were sealed and incubated at 30°C and visually evaluated for growth after three to five days once growth was visible in the growth control wells. Azithromycin and clarithromycin conditions were reassessed after 14 days of incubation to evaluate inducible macrolide resistance. MICs were recorded as the lowest drug concentration preventing visible mycobacterial growth. For further analyses, we used either log2 transformed MIC values or applied previously reported *M. abscessus* resistance breakpoints (clarithromycin MIC ≥ 8 µg/ml; amikacin MIC ≥ 64 µg/ml (*7*)).

*M. tuberculosis* drug susceptibility testing was done analogous to *M. abscessus*, except *M. tuberculosis* isolates were cultured in 5 ml of Middlebrook 7H9 broth, supplemented with 0.4 % glycerol, 10 % OADC and 0.05 % Tween 80 in 50 ml tubes (*M. tuberculosis* $mc^2$7000 was additionally supplemented with 100 µg/ml pantothenate) and that MIC plates were assessed after 14 days. Bedaquiline, clofazimine, delamanid, ethambutol, linezolid, moxifloxacin, pretomanid, rifampicin, rifapentin, SQ109 and sutezolid were assessed in log2-fold concentration steps between 2 ng/ml – 64 µg/ml, whereas isoniazid and pyrazinamide were assessed between 4 ng/ml – 128 µg/ml.

##### ***In vivo*** **data**

The *in vivo* classifications of *M. tuberculosis* drug regimens “as good or worse than standard-of-care (SOC; where isoniazid-rifampicin-pyrazinamide-ethambutol or isoniazid-rifampicin-pyrazinamide were considered SOC)” or “better than standard-of-care” were obtained from two previous studies. *In vivo* classifications based on mouse models were obtained from (*8*), and classifications based on phase 2a and 2b clinical trials were obtained from (*9*). All similar to or better than SOC drug combinations from the relapsing mouse model (n = 46) and clinical studies (n = 14) were assessed with ASCT. These drug combinations and classifications of single drugs were also used to evaluate the performance of ASCT in bactericidal mouse models (common mouse strains: n=48; C3HeB/FeJ mouse strain: n=15). Clinical metadata of patients with respiratory *M. abscessus* infection were available for a subset of patients. Treatment outcomes were assessed in patients meeting the ATS/IDSA criteria of NTM pulmonary disease (*10*). Patients with positive cultures after six months of *M. abscessus* treatment were considered as persisting infections, and patients with microbiological clearance as cleared infections. Growing isolates were excluded from the analysis of bacterial killing. The discriminative performance for predicting the success of drug regimens in mouse or patient populations (*M. tuberculosis*) and the clinical outcome in individual patients (*M. abscessus*) was assessed with logistic regression. The discriminative performance was quantified with the area under the receiver operating curve (AUC-ROC). Fifty curves were generated for each condition by randomly selecting 80 % of the samples. ROC curves were compared using DeLong's test.

#### Antimicrobial Single-Cell Testing

##### *ASCT - experimental setup*

*M. abscessus* and *M. tuberculosis* isolates were cultured in 5 ml of Middlebrook 7H9 broth, supplemented with 0.4 % glycerol, 10 % OADC and 0.05 % Tween 80 in 50 ml tubes (*M. tuberculosis* $mc^2$7000 was additionally supplemented with 100 µg/ml pantothenate). The cultures were incubated at 37°C with shaking at 150 rpm until mid-log phase (McFarland: 5-8). In *M. tuberculosis*, we also assessed two starvation conditions: Phosphate Buffered Saline (PBS)-starvation and pantothenate-starvation. For PBS-starvation, mid-log phase bacteria (H37Ra or *M. tuberculosis* $mc^2$7000) were washed and resuspended in PBS, the 50 ml tube filled with PBS and incubated for 14 days at 37°C without shaking. For pantothenate-starvation, *M. tuberculosis* $mc^2$7000 (H37Rv ΔpanCD ΔRD1) was incubated for 14 days with Middlebrook 7H9, supplemented with 0.4 % glycerol, but not pantothenate.

After growth or starvation, mycobacteria were centrifuged at 3000 g for 10 min, and the resulting pellet was resuspended in Middlebrook 7H9 (for growth conditions and pantothenate-starvation), in Middlebrook 7H9 with pantothenate supplementation (for growth conditions in *M. tuberculosis* $mc^2$7000) or in PBS (for PBS-starvation). To achieve single-cell suspensions, large clumps were removed by low-speed centrifugation (200 g, 3 min), followed by serial filtration of the supernatant through 5 µm and 1.2 µm filters (Sartorius Minisart). In every sample, bacterial densities were assessed using $OD_{565}$ measurements, and the live-cell fraction was quantified using propidium iodide staining and imaging. Bacterial samples with a live-cell fraction below 95 % were discarded and reprocessed. Single-cell suspensions were immediately used (*M. tuberculosis*) or frozen at -80°C (*M. abscessus* clinical isolates).

Antimicrobial Single-Cell Testing (ASCT) uses a dual-layer approach: the first layer consists of an agar pad containing and immobilising bacteria, and the second layer comprises drug-containing solutions (***Fig. 1A***). To prepare the agar pad, ultra-low gelling temperature agarose (ULGA; Lonza SeaPrep) and Middlebrook 7H9 were dissolved in hot dH2O. Additional dissolution was facilitated through heating, where any evaporated water was replaced to maintain constant 7H9 and agarose concentrations. Once the solution cooled to 50°C or lower, OADC and glycerol were added. The mixture was then filtrated through a 0.22 µm filter and propidium iodide was added. This agarose solution was aliquoted as needed. The final gel pad solution contained 0.4 % agarose, 1x Middlebrook 7H9, 10 % OADC, 0.4 % glycerol, 8 µg/ml propidium iodide and approximately $5*10^6$ bacteria/ml. For non-starving *M. tuberculosis* $mc^2$7000, the agar pad also contained 100 µg/ml pantothenate. In pantothenate-starvation conditions, the gel pad did not contain pantothenate, and in PBS-starvation media and all supplements were replaced with 1x PBS.

All preparations were made in 96 well-plates using a 96-channel pipette (Mini96, Integra Biosciences). 7 µl of the isolate-agar-PI solution was dispensed using liquid handling (I.DOT, Dispendix) into each well of a 1536-well plate (Greiner, Screenstar). At least one column and row were left as buffer wells at the edges. All materials used in the agarose preparation, including chemicals, pipette tips, filters, the plate, etc., were preheated to 37°C. The 1536-well plate was centrifuged at 37°C and 3,000 g for 30 min to position the bacteria at the bottom of the wells for imaging. This step was followed by cold centrifugation at 4°C and 1,500 g for 20 min to solidify the ULGA. After leaving the plate for 30 min at room temperature, a defined drug solution (4 µl) dissolved in Middlebrook 7H9 or PBS (for PBS-starvation) was added to each well. The plate was then sealed with parafilm, covered with a lid, and remained for another 60 min at room temperature before being placed into the microscope's pre-heated live-cell chamber (37°C; Life Imaging Services). To evaluate the impact of gel pad volumes and concentrations on antibiotic killing, we tested various volumes (4, 5, 6, 7 µl) and concentrations (0.3, 0.4 and 0.5%; ***fig. S1G-H***).

Images were captured with a Nikon Ti2-E inverted microscope using a CFI Plan Apo Lambda 40x NA 0.95 or a CFI Plan Apo Lambda 100x NA 1.45, and the Perfect Focus System for maintenance of the focus over time. PI fluorescence was excited with a Spectra III Light Engine (Lumencor) at 555 nm and collected with a penta-edge 408/504/581/667/762 dichroic beam splitter and a 440/40, 520/21, 606/34, 694/34, 809/81 penta-bandpass filter. 40x 12-bit and 100x (optionally with 1.5x zoom lens) 16-bit images were acquired with a Photometrics Kinetics camera, controlled with the Nikon NIS acquisition software. For 40x, the fluorescence excitation light intensity was set to 10% and the camera exposure time to 20 ms, and the brightfield illumination light intensity was set to 50% and the camera exposure time to 8 ms. These settings were kept the same for all experiments and conditions for comparability. For 100x acquisition, the excitation light intensity and camera exposure were adjusted accordingly.

The NIS JOBS module was used for image acquisition automation. Briefly, 9 equally spaced fields of view for each well and imaging time point were acquired. For *M. abscessus*, images were captured every 2.5 h for a total duration of 72 h and 29 time points; for *M. tuberculosis*, images were acquired every 4 h for a duration of 168 h (42 time points in total). Imaging one time point took approximately 130 min (1260 wells with 9 fields of view per well). At the end of image acquisition, imaging data (*M.* abscessus: up to 11,340 movies and 7.8 TB per experiment; *M. tuberculosis*: up to 11 TB per experiment) were transferred to the high-performance computing cluster at the University of Basel (sciCORE) for subsequent image and data analysis.

To evaluate ASCT-based time-kill kinetics of *M. tuberculosis,* we exposed *M. tuberculosis* to previously published drug combination regimens (*8*, *9*) at the maximum blood concentration ($C_{max}$) achievable during therapeutic dosing in humans (9 replicates for each condition): bedaquiline 1.1 µg/ml, clofazimine 1.25 µg/ml, delamanid 0.5 µg/ml, ethambutol 4 µg/ml, isoniazid 4.5 µg/ml, linezolid 19 µg/ml, moxifloxacin 4 µg/ml, pretomanid 7.8 µg/ml, pyrazinamide 40 µg/ml, rifampicin 16 µg/ml, rifapentin 19 µg/ml, SQ109 0.026 µg/ml, sutezolid 1.48 µg/ml.

For drug tolerance profiling, all clinical *M. abscessus* isolates were assessed within a single ASCT plate for a single antibiotic condition in triplicate. The drug concentrations used were based on the MICs for ATCC-19977, determined similar to CLSI criteria (*7*). In contrast to CLSI, they were assessed in 7H9 supplemented with 0.4 % glycerol, 10 % OADC and 0.05 % Tween 80 and quantified with optical densities ($OD_{600}$). Growth inhibition at increasing drug concentrations was fitted with a four-parameter log-logistic model using the R drc package (*11*). The MIC was defined as the drug concentration that prevented 90 % growth ($IC_{90}$). Antibiotic time-kill kinetics for all eight drugs were evaluated at two different drug concentrations: a lower concentration (10-fold the MIC of ATCC-19977) and a higher concentration (20-fold the MIC of ATCC-19977). The specific drug concentrations (low and high) tested were as follows: amikacin 26 and 52.1 µg/ml, azithromycin 63.5 and 127 µg/ml, cefoxitin 63.1 and 126.2 µg/ml, imipenem 37.7 and 75.4 µg/ml, minocycline

67.2 and 134.4 µg/ml, moxifloxacin 8.5 and 17 µg/ml, linezolid 38.5 and 77.1 µg/ml, tigecycline 27.2 and 54.3 µg/ml.

##### *ASCT - image analyses*

Image processing of every image time-lapse (all imaging time points of a single field) consists of five individual image analysis steps: background control, cell segmentation, cell classification, drift correction, and cell tracking.

To increase the accuracy of fluorescence intensity quantifications, we applied BaSiC, a method that automates background correction through low-rank and sparse decomposition via Fiji (*12*, *13*). This method adjusts for spatial and temporal variations in background fluorescence, which commonly arise from uneven and repetitive illumination. Employing the default BaSiC settings, we achieved consistent fluorescence signals across different conditions, well positions and time (***fig. S1B***).

Bacteria were segmented using a combined pixel and object classification approach implemented in ilastik (*14*). In the pixel classification task, each pixel was subjected to supervised Random Forest classifiers (100 trees) to differentiate between the background and cellular structures. Within the object classification task, the pixel classification map, estimating the probability of each pixel belonging to the background or cellular class, was smoothed and thresholds were applied to generate object features such as intensity statistics and shape descriptors. These features were used to manually train the following object classifiers: PI+ single cells, PI- single cells and clumps. Pixel and object classifiers were manually trained on approximately 30 time-lapses, including different *M. abscessus* strains and antibiotic conditions. These classifiers were then applied to all data (over 200,000 time-lapses) in batch mode. To assess the fate of PI+ and PI- bacteria upon antibiotic washout, we trained bacteria with ambiguous PI signals as an additional object class. This class was excluded from single-cell growth assessments. To validate segmentation accuracy, three randomly selected time-lapse datasets were manually annotated (with automated support) to generate ground truth pixel prediction maps. The same time-lapses were then processed by ASCT pixel classification. Ground truth and ASCT pixel classification was compared using the ImageJ CLIJ2 plugin (*13*). Segmentation accuracy (i.e., object overlap) was quantified with the Jaccard Index (***fig. S1A***). To quantify PI classification accuracy, 1,600 PI positive and 1,600 PI negative single cells were manually annotated within five timelapse datasets, including different *M. abscessus* isolates and treatment conditions. The ASCT PI classification algorithm was than compared with manually annotated objects using the R caret package (***fig. S1C***) (*15*).

To optimise downstream analysis and especially bacterial tracking, we corrected for any drift in imaging fields, which mainly occurred at the beginning of each experiment due to thermal changes (thermal drift). The drift of fields was determined by identifying the maximum cross-correlation between two consecutive images by shifting the subsequent image along the x and y coordinates. Drift correction for large-scale time-lapse data is computationally expensive. Therefore, we used smaller field sections (600 x 600 pixels rather than the 2400 x 2400 pixels of the whole field) and adapted the process to distinct imaging time points (frames), given that the drifts were minor after the initial imaging frames. We extracted a window of 600 x 600 pixels from the binary map of segmented objects in frame 0 (centred at $c_1$ = (800,800)), the very first image of a time-lapse. We also extracted a window of 600 x 600 pixels from frame 2 and shifted this section, one pixel at a time, up to 200 pixels away from $c_1$ in each direction, creating 401 x 401 windows. The windows from frame 1 were overlaid on the frame 0 window to find maximum alignment. Given that cells are not always uniformly distributed or can lose focus, the same strategy was performed with two other windows ($c_2$ = (1300,1300) and $c_2$ = (1700,1700)). The maximum of the cross-correlation across the three locations was used. From frame 3 onward, windows were shifted by fewer pixels to achieve a 75 % alignment likelihood between the frames. The shifts were initially set to 25 pixels and consecutively adjusted to 25, 50, 100 and 200 pixels if the alignment likelihood was not achieved.

To automatically follow the behaviours of individual cells during the experiment, we established a custom script to track segmented objects across imaging frames. For each frame, we extracted object features from the segmentation output obtained via brightfield imaging using the MATLAB function *regionprops*. Our cell-tracking algorithm aims to link corresponding objects between the initial frame 0 and subsequent frames, with $n_t$ denoting an object $n$ at time frame $t$. Object homology $H$ between frames was determined by comparing features of individual objects. A graph of linking objects ($n_t$, $n_0$) at frame $t$ was represented by a 3-dimensonal tensor $H(n_0, n_0+N, t)$, where $N_0$ is the population of objects at frame 0. The $H$ tensors at each frame were composed of dissociation weights based on the following features: 1) the distances between centroids of objects $n_t$ and $n_0$; 2) the changes in the area of objects; 3) changes in object orientation; and 4) changes in the convex hull area of the object. We assigned higher weight to distances (raised to the power of 4) and a weight of one to the other features.

$$H(n_0, n_t, t) = Dist\big(cent(n_t), cent(n_0)\big)^4 \frac{\left|Area(n_0) - \frac{Area(n_t)}{dA/dt}\right|}{Area(n_0)} \frac{\left|CoHull(n_0) - \frac{CoHull(n_t)}{dA/dt}\right|}{CoHull(n_0)} ang(n_0, n_t)$$

All possible homology values were integrated into an object matrix. Rather than comparing each frames object to find the closest match, we employed a comprehensive local minimum strategy. The pair of two objects with the highest homology (lowest values in the matrix) was identified as a link in an array and subsequently removed from the matrix. This process was iteratively repeated to identify and remove further links until a homology threshold of 1 x $10^8$ was reached. This local minimum strategy allowed individual objects to be linked throughout the time-lapse to generate single-cell trajectories. Objects of abnormal size, merging objects, moving objects, objects that increased or decreased in object area, objects with swapping labels, or objects that were not tracked for more than two frames within the time-lapse were discarded. Wells with less than 1000 tracked bacteria for *M. abscessus* and less than 500 tracked bacteria for *M. tuberculosis* were discarded. The time of PI positivity of a single cell was defined as the first PI+ frame of two consecutive PI+ frames.

##### *ASCT - data analyses*

After quantifying the morphology and viability trajectories of each bacterium, the imaging data were further analysed. Bacterial growth in ASCT was defined as an increase in total object area greater than 3.1-fold during antibiotic exposure. This threshold was first defined by comparing growing and non-growing cells and then validated in *M. abscessus* isolates with and without inducible macrolide resistance during azithromycin exposure (***fig. S1D***). If growth was detected in two third or more of the replicates for a given isolate-drug condition, the condition was considered growing and removed from the killing analyses.

The reproducibility of *M. abscessus* time-kill kinetics was assessed by comparing live-cell fractions at distinct time points (3, 6, 9, 12, 24, 36, 48, 60 and 72 hours). Exact live-cell fractions at given time points were interpolated from overall time-kill kinetics. The coefficient of variation (CoV) was calculated for each triplicate at every time point. All triplicates were included in downstream analyses if the mean CoV of all three replicates fell within 3 SD of the overall mean triplicate CoV distribution per drug condition. Triplicates exceeding this threshold were reduced to the best-performing duplicate. These duplicates were retained if their mean duplicate CoV was within 3 SD of the overall mean duplicate CoV per drug condition; otherwise, the isolate-drug pair was excluded from further analyses. A minimum of two reproducible time-kill curves per isolate-drug condition were required for downstream analyses of *M. abscessus* kill analyses. To assess for outliers in our *M. abscessus* drug tolerance assessment, we performed principal component analyses for every antibiotic condition based on live-cell fractions at 3, 6, 9, 12, 24, 36, 48, 60 and 72 hours. Isolate-drug conditions with a distance of more than 3 SD from the mean, assessed with Mahalanobis distance and calculated using the first two principal components, were excluded from further analyses. In total, 957 (14.7 %) *M. abscessus* isolate-drug pairs were excluded from time-kill analysis due to bacterial growth (mostly azithromycin and minocycline) and 156 (2.4 %) due to poor reproducibility or being outliers. Due to the larger number of replicates and small number of isolates, the strategies to account for reproducibility and outliers were not applied to *M. tuberculosis*.

Imaging data were acquired consecutively for each well at different time points. The live-cell fraction at 0 hours (time point 0) was extrapolated by fitting a logistic function to the live-cell fractions from the first 5 imaging frames, corresponding to approximately 13 hours (all conditions except imipenem and cefoxitin). Due to the rapid killing observed in imipenem and cefoxitin conditions, resulting in inaccurate fitting, the maximum live-cell fraction observed for each isolate in any of the other drug conditions was used as the isolate-specific live-cell fraction at 0 hours. For *M. abscessus* isolates with a live-cell fraction below 80% and isolates which had less than 1,000 cells per well across multiple wells were excluded from further analyses. Using the 0-hour live-cell fraction, all frames were normalised to an initial live-cell fraction of 100%. Overall, antibiotic killing was quantified as the arithmetic mean of the area under the 72-hour (*M. abscessus)* or 168-hour (*M. tuberculosis)* time-kill curve, ranging from 1 to 0 (maximum survival to most rapid killing, respectively).

**Drug tolerance analyses**

We assessed the relationship between *M. abscessus* drug tolerance phenotypes, growth rates, lag times and MICs using Pearson correlation and $R^2$ values. Spearman correlation was employed to examine the contribution of outliers. For drug clustering in drug tolerance space, a Spearman correlation matrix was generated based on pairwise comparisons of the area under the time-kill curves from 350 clinical *M. abscessus* isolates. Only isolates with four or less missing drug tolerance values (out of 16) were used. Principal component analysis was applied to the correlation matrix to visualise drug clustering.

**Single-cell growth assessment**

We also applied the tracking algorithm to identify whether single cells are able to grow (form microcolonies) after antibiotic washout. To achieve tracking accuracy and computational efficacy, we employed a backward approach. We identified and categorised objects with a size 5 – 15 times the median size of single cells as microcolonies and recorded the area, appearance time, and centroids. Analogous to the homology matrix to track single non-growing cells, we used the homology index to track microcolonies backwards to the final single-cell object, i.e., the initial originating cell. We focused on two features: the object area, which is important particularly to track large objects, which grow and potentially move; and centroids, which are critical to track single non-moving cells. With this approach, we could determine whether specific objects generated microcolonies.

##### Whole-genome sequencing

*M. abscessus* isolates were cultured on solid media and colony sweeps were collected (*2*, *16*). DNA extraction was performed using the Qiagen QIAamp DNA mini kit. DNA libraries were constructed with unique identifiers for each isolate and sequenced using multiplexed paired-end sequencing. *De novo* genome assemblies were assessed for quality. Assemblies with a length longer than 6 Mb, more than 300 contigs, an average depth below 30 x, a coverage of the reference genome below 50 % or presumed mixed infection were discarded. Sequence reads were mapped to the *M. abscessus* ATCC-19977 genome using BWA, followed by INDEL realignment (*17*, *18*). Single nucleotide polymorphisms (SNPs) and small insertions/deletions (INDELs) were identified using bcftools and annotated with SNPeff (*19*, *20*). SNPs were filtered to require a minimum base call quality of 50, a minimum mapping quality of 20, and at least 8 matching reads covering a SNP (3 per strand). To assess larger deletions, ATCC-19977 was partitioned into regions of 20 bp with 10 bp overlaps (*21*). The coverage of these regions in clinical isolates was assessed with sambamba (*22*). Large deletions were defined as two consecutive windows with a mean coverage of 5 x or below, occurring in at least 5 % of all genomes. Variants with identical distributions were collapsed into a single variant. Maximum likelihood trees were generated using FastTree, inferred from core SNPs and visualised with iTOL (*23*, *24*). Mutational aminoglycoside resistance was evaluated with mutations in the *rrs* genotype and macrolide resistance with mutations in *rrl* (*25*, *26*).

##### Heritability estimations

Unitigs for all isolates were extracted using the unitig-caller tool, which utilises an FM-index built around the Bifrost API (*27*, *28*). A similarity matrix was generated from phylogenetic distances, which was then used to correct for population structure. FaST linear mixed models were used to estimate narrow sense heritability ($h^2$), representing the proportion of variance in the phenotype attributable to genetic variation (*29*, *30*). Random chance was assessed by shuffling each drug tolerance phenotype across *M. abscessus* isolates and calculating heritability (10 times for each drug tolerance phenotype; 160 times in total).

##### Phenogenomic analysis

We performed genome-wide associations studies (GWAS) to analyse approximately 300,000 *M. abscessus* genetic variants, including SNPs, INDELs and large deletions, in relation to drug tolerance phenotypes. Variants were classified by presumed genetic effects: low effect (intergenic variants, synonymous SNPs), moderate effect (non-synonymous SNPs, inframe INDELs) and high effect (frameshift variants, start/stop alterations, large deletions). We applied linear mixed models to account for population structure, integrating a relatedness matrix, and quantified associations with the Wald test (*21*, *31*). We used a Bonferroni threshold of $1.7*10^{-7}$ to control for multiple hypothesis testing. To summarise GWAS hits, we extracted the top five genes showing the strongest association per phenotype (for moderate or high effect variants). The top associations of these genes with all tolerance phenotypes were then shown in a heatmap (moderate or high effect variants). Associations were plotted using LocusZoom (*32*).

##### Generation of *the MAB_0233* knockout mutant

*MAB_0233* knockout mutants (Δ*MAB_0233*) were generated on the *M. abscessus* ATCC-19977 background using ORBIT (*33*). pKM444 (RecT-Int-expressing plasmid, $Kan^r$) and pKM496 (for gene deletion by integration into target gene, $Zeo^r$) was a gift from Kenan Murphy (Addgene plasmid #108319 and #109301) (*33*). pMV261 (replicative vector for gene complementation) was obtained from NovoPro Biosciences (*34*). To amplify plasmids, *Escherichia coli* DH5α cells were grown on LB agar plates supplemented with 50μg/ml kanamycin (pKM444 or pMV261) or 50μg/ml zeocin (pKM496).

Liquid cultures of *M. abscessus* ATCC-19977 were incubated at 37°C, 150 rpm for two days. Cells were diluted to an $OD_{600}$ of 0.03 in 20 ml of Middlebrook 7H9, supplemented with 0.4 % glycerol, 10 % OADC and 0.05 % Tween 80 in a 250 ml baffled flask. The culture was placed at 30°C, 100 rpm for 18 to 20 hours. The culture was then washed three times with 20 ml ice-cold sterile 10 % glycerol supplemented with 0.05 % Tween 80 (washing solution). Following the third wash, the cells were collected by centrifugation and resuspended in 200 μl of washing solution. The ORBIT plasmid pKM444 (200 ng) was mixed with competent *M. abscessus* and rested on ice for 5 min. The cells were electroporated (2.5 kV, 1,000 Ω, and 25 μF) using a 2 mm gap width electroporation cuvette. Following electroporation, the cells were resuspended in 1.5 ml Middlebrook 7H9 medium and placed at 30°C, 100 rpm for 18 to 20 h. The recovered cells were pelleted at 3000 g (25°C, 5 min), resuspended in 100 μl Middlebrook 7H9 medium, and plated on 7H11 plates supplemented with 0.5 % glycerol, 10 % OADC and 250 μg/ml kanamycin. The plates were incubated at 30°C for 3-4 days.

Colonies resulting from pKM444 transformation were picked and grown on Middlebrook 7H11 agar plates supplemented with 0.5 % glycerol, 10 % OADC and 250 μg/ml kanamycin. Cells were harvested by centrifugation and resuspended in TRIzol (Thermo Fisher). Cell disruption was performed by bead-beating using Lysing Matrix B beats (200 μl) in a FastPrep-24™ Classic instrument at 6.5 m/s for 60 s followed by 60 s incubation on ice. This process was repeated for three cycles. Total DNA was extracted using the DNA miniprep kit (Zymo Research). The presence of the pKM444 plasmid was verified by PCR, using DreamTaq™ DNA Polymerase (Thermo Fisher Scientific). Primers specific to the plasmid (forward: CACGTTGTGTC-TCAAAATCTC; reverse: CGATAACGTTCTCGGCTC) were used to amplify a target segment of the plasmid. PCR products were resolved by agarose gel electrophoresis, and bands of the expected size were excised and submitted for Sanger sequencing to verify plasmid presence.

The oligonucleotide sequence consists of the 48 bp Bxb1 *attP* site (or the reverse complement) flanked by 60 bp upstream of the initiation codon and 60 bp downstream of the stop codon (*33*). The targeting oligonucleotide (sequence for *MAB_0233*: CGGCCACATGTTTTGTGCCGCTAGGG-GAAATCAGCTCGGCATCGTCCGTGTGCCGTTGTTGGTTTGTACCGTACACCACTGAGACCGC GGTGGTTGACCAGACAAACCCACATACCCTGATGCGAGTTCAACTGCCATCTGTTGCCCCCTT CGGGCAATCGGTGCAGC) was acquired from IDT with 4 nmole Ultramer™ DNA Oligo property and diluted in nuclease-free water to 1 μg/μl.

*M. abscessus* ATCC-19977, harbouring the pKM444 plasmid, was cultured in Middlebrook 7H9 medium with 5 μg/ml anhydrotetracycline (ATc). A 20 ml culture was prepared in a 250 ml baffled flask wrapped in aluminium, washed as described above and resuspended in 200 μl washing solution. 2 µl of 1 µg/µl oligonucleotide, guiding the payload plasmid pKM496 to the target gene, was aliquoted into 1.5 ml microcentrifuge tubes and denatured at 95°C for 5 min to prevent secondary structure formation. The denatured oligonucleotides were cooled on ice for at least 5 min before adding 200 ng pKM496 (*33*). Subsequently, 200 µl of cells were added to the tube, gently mixed by pipetting, and incubated for 5 min. Electroporation, recovery, and plating were performed as described above, except recovered cells were plated on 7H11 plates supplemented with 100µg/ml zeocin. DNA extraction and PCR were performed as described using primers flanking *MAB_0233* and pKM496 (forward: CGCTCACAACTGAATACCC; reverse: CCTGGTATCTTTATAGTCCTGTC).

**Gene complementation**

To complement Δ*MAB_0233*, the gene was amplified by PCR and cloned into pMV261 downstream of the mycobacterial strong constitutive promoter, *pHSP60* (*34*). In addition, the T2 terminating region from *E. coli rrnB* was added to the ORF's end as an efficient terminator of transcription. To eliminate the pKM444 plasmid, which confers kanamycin resistance, the *M. abscessus* knockout strain Δ*MAB_0233* was subjected to serial dilutions and growth cycles until cured. Cured strains were transformed with pMV261 following the same transformation protocol as described for pKM444.

As control MAB-ATCC-pKM444 was used. The RNA of log phase cultures of the control strain, Δ*MAB_0233* and Δ*MAB_0233*::*MAB_0233* was isolated using the RNA miniprep kit (Zymo Research). cDNA was prepared using the High-Capacity cDNA Reverse Transcription Kit (Zymo Research). cDNA levels were quantified by quantitative real-time PCR (qRT-PCR) on an Applied Biosystems qPCR machine using a PowerUp SYBR Green Master Mix (Thermo Fisher Scientific) and analysed by the ΔΔCt method. Gene expression was controlled using the housekeeping genes *MAB_3009* (sigA) and *MAB_ 3869c*. Two sets of primers covering approximately 150 bp sequences of the beginning and the end of the *MAB_0233* ORF were designed (primer_1_FWD: GCGAAGCCTTCGCCAAAGCTC, primer_1_REV: CCTCGTTGAGCTTTTCCAGCGC, primer_2_FWD: CGCAACGTCTCCGATGGGAAACC, primer_2_REV: GAGTTGAGCGGCGTCCATGC).

#### SUPPLEMENTARY FIGURES

### Supplementary Figure 1

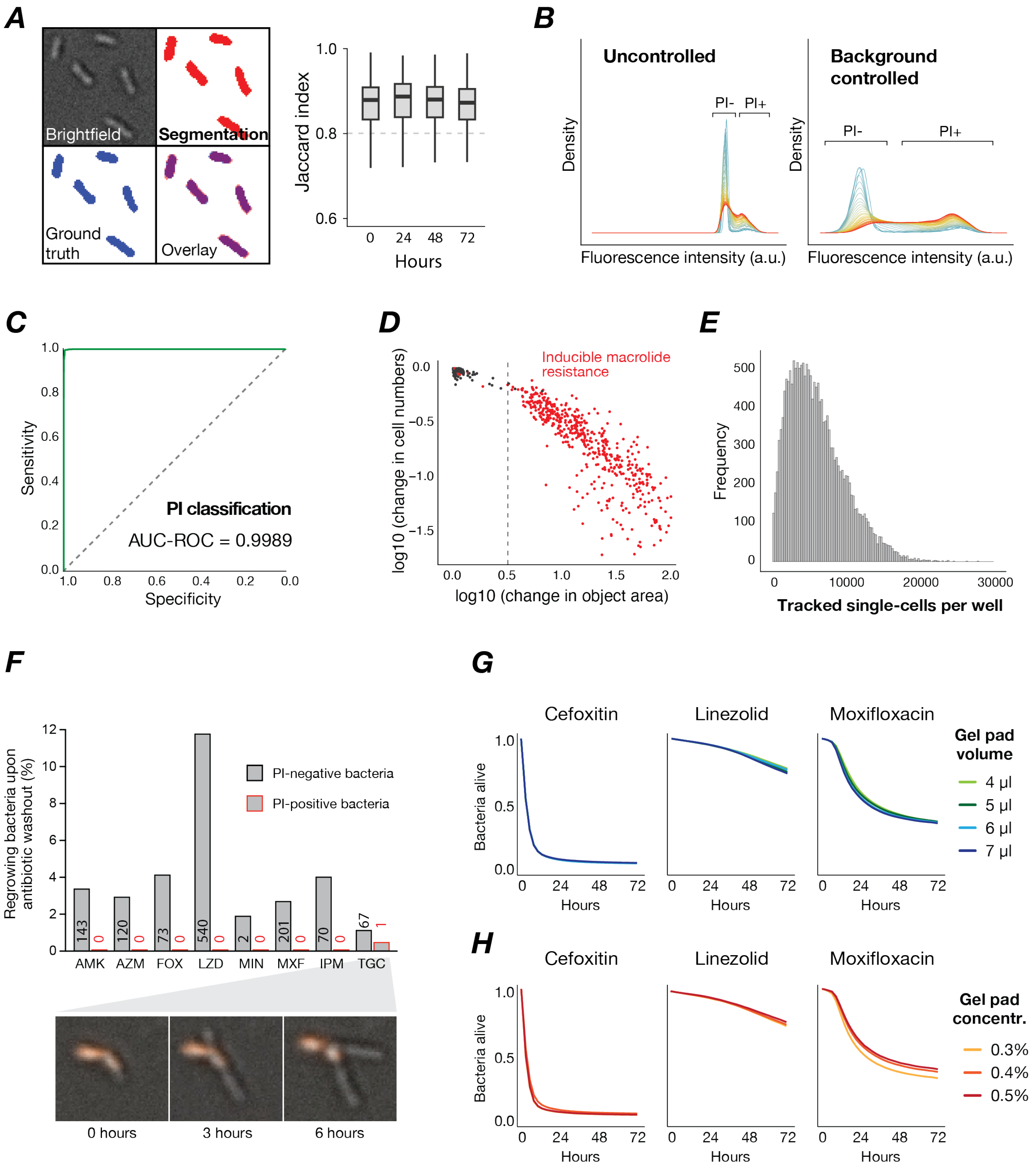

**Figure S1. Antimicrobial Single-Cell Testing.**

**(A)** Overlay of manually annotated “ground truth” bacterial segmentation with automated object segmentation (pixel classification) of *M. abscessus* brightfield images. The Jaccard index quantifies overlay accuracy across imaging time points during time-lapse acquisition of 5,887 bacteria. The dashed line indicates high segmentation accuracy. **(B)** Changes in dynamic range following BaSiC background correction. Mean PI values per single *M. abscessus* bacterium before and after correction over 72-hour time-lapses (first imaging time point: blue, last time point: red) shown in density plots. **(C)** Accuracy of automated PI classification in predicting “ground-truth” classes of PI-positive and PI-negative single cells. **(D)** Changes in cell count and object area (assessed with ASCT) over 72 hours under azithromycin treatment across 406 *M. abscessus* isolates. Isolates with (red) and without (black) inducible macrolide resistance are highlighted. The dotted line indicates the ASCT growth threshold. **(E)** Number of single *M. abscessus* bacteria tracked over 72 hours per well within a 1536-plates. Data from 406 *M. abscessus* isolates, and about 130 million tracked bacteria across eight drugs and two concentrations (data from Figure 3). **(F)** Fraction of bacteria (including numbers) showing regrowth after 24-hour antibiotic treatment and antibiotic washout across PI-negative and PI-positive bacteria. The single regrowing PI-positive bacterium is highlighted in the insert. **(G)** Effect of different gel pad volumes on *M. abscessus* time-kill kinetics. **(H)** Effect of different gel pad agarose concentrations on *M. abscessus* time-kill kinetics.

### Supplementary Figure 2

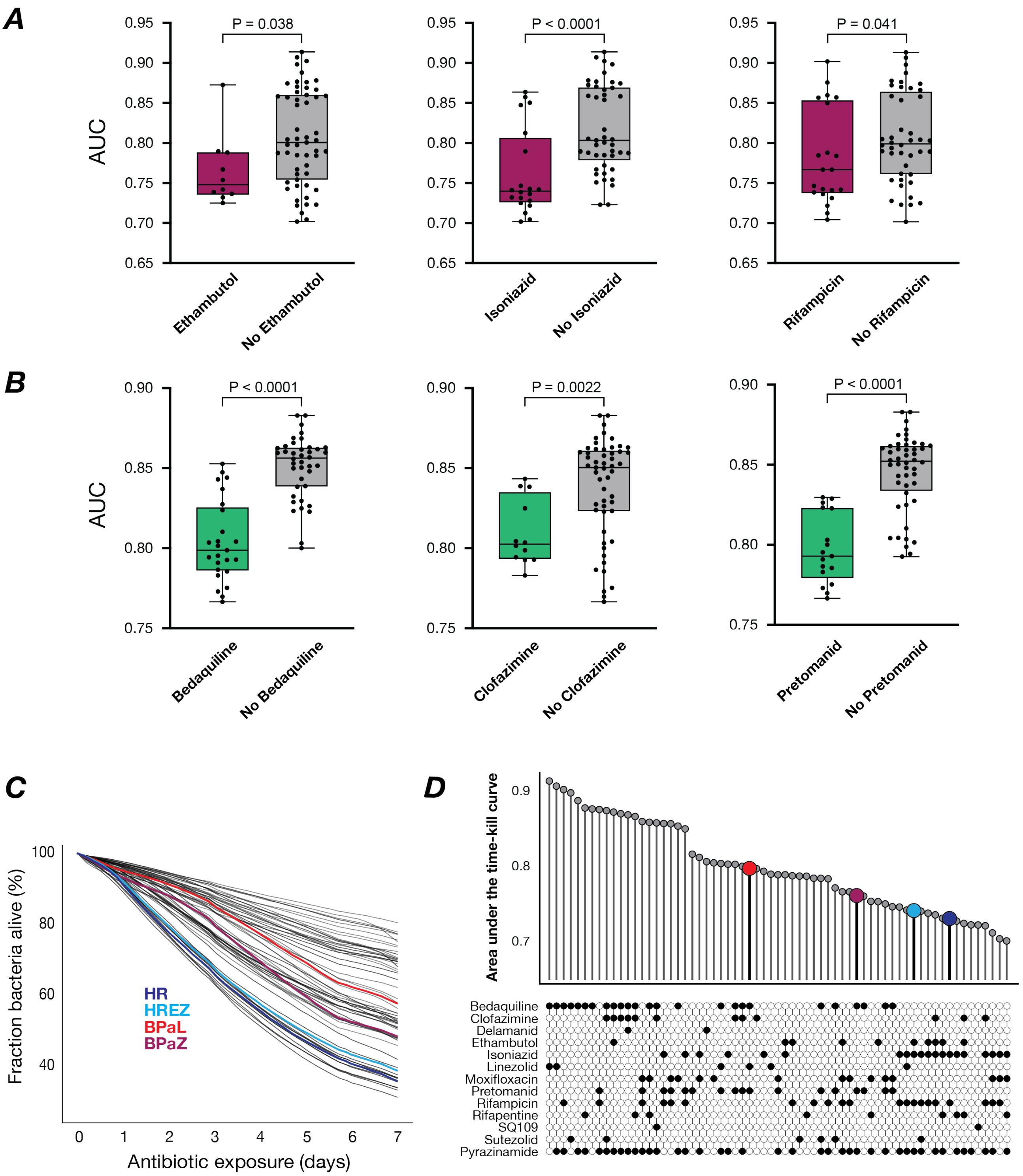


**Figure S2.** ***M. tuberculosis*** **time-kill kinetics (A-B).** Time-kill curves (AUCs) of *M. tuberculosis* drug combinations were assessed in regard to the presence and absence of individual drugs. Three drugs demonstrated different killing upon **(A)** exponential growth (averaged from mc$^2$7000 and H37Ra strains), whereas other three drugs showed different killing in **(B)** starvation conditions (averaged from three starvation conditions). Groups were compared using the Mann-Whitney U test. Each dot indicates a drug regimen. Boxplots show the median, interquartile range and total range of AUC values. **(C)** ASCT-based time-kill kinetics of exponentially growing *M. tuberculosis* (H37Ra) exposed to 65 drug regimens, with the following regimens highlighted: isoniazid-rifampicin (HR; dark blue), isoniazid-rifampicin-ethambutol-pyrazinamide (HREZ; light blue), bedaquiline-pretomanid-linezolid (BPaL; red), and bedaquiline-pretomanid-pyrazinamide (BPaZ; purple). **(D)** Mean area under the kill curve for *M. tuberculosis* drug regimens under exponential growth averaged across the two *M. tuberculosis* strains.

### Supplementary Figure 3

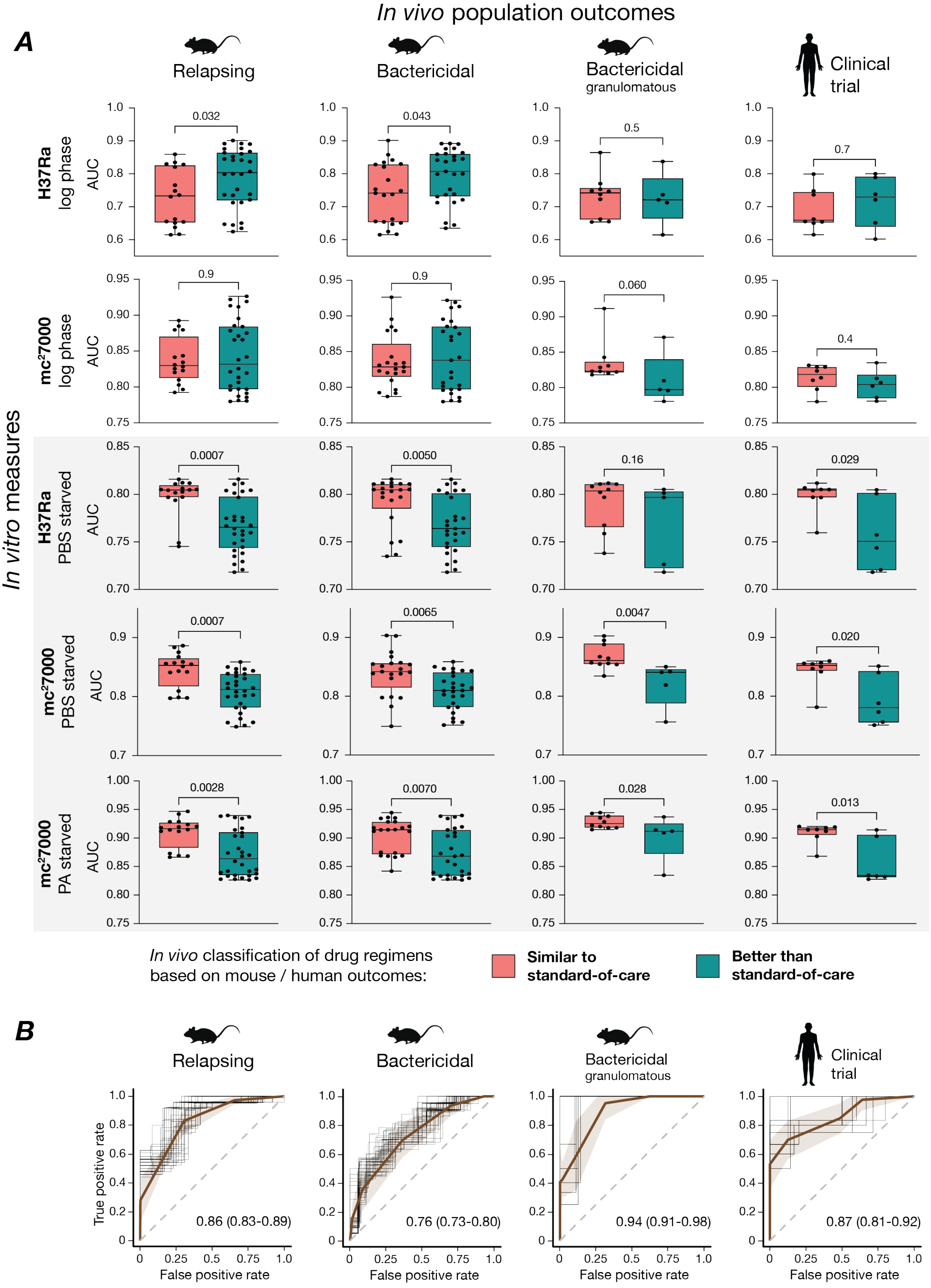

**Figure S3. Association of *in vitro* killing with *in vivo* outcomes of *M. tuberculosis* drug regimens. (A)** *M. tuberculosis* drug regimens were previously classified as similar or better than standard of care (SOC) based on the performance in relapsing mouse models (RMM), bactericidal mouse models (BMM) of common mouse strains, the granulomatous C3HeB/FeJ strain and clinical studies (*8*, *9*). Time-kill curves (AUCs) upon exponential growth and starvation conditions of two *M. tuberculosis* isolates were used to compare “similar to SOC” versus “better than SOC” classifications using the Mann-Whitney U test (P indicated). Each dot represents a drug regimen. Boxplots show the median, interquartile range and total range of AUC values. **(B)** Performance of time-kill kinetics (AUC averaged across three *M. tuberculosis* starvation models) for predicting *in vivo M. tuberculosis* outcomes (“similar to SOC” versus “better than SOC” drug regimens). Using logistic regression, fifty ROC curves were generated for every condition by randomly selecting 80% of the samples. The brown line represents the mean receiver operating characteristic (ROC) curve, the shaded area one standard deviation. Mean area under the ROC curves and 95% confidence intervals are presented. (RMM: n = 46, BMM common strains: n = 48, BMM C3HeB/FeJ: n = 15, Clinical bactericidal activity: n = 14). PA indicates pantothenate, PBS phosphate-buffered saline.

#### Supplementary Figure 4

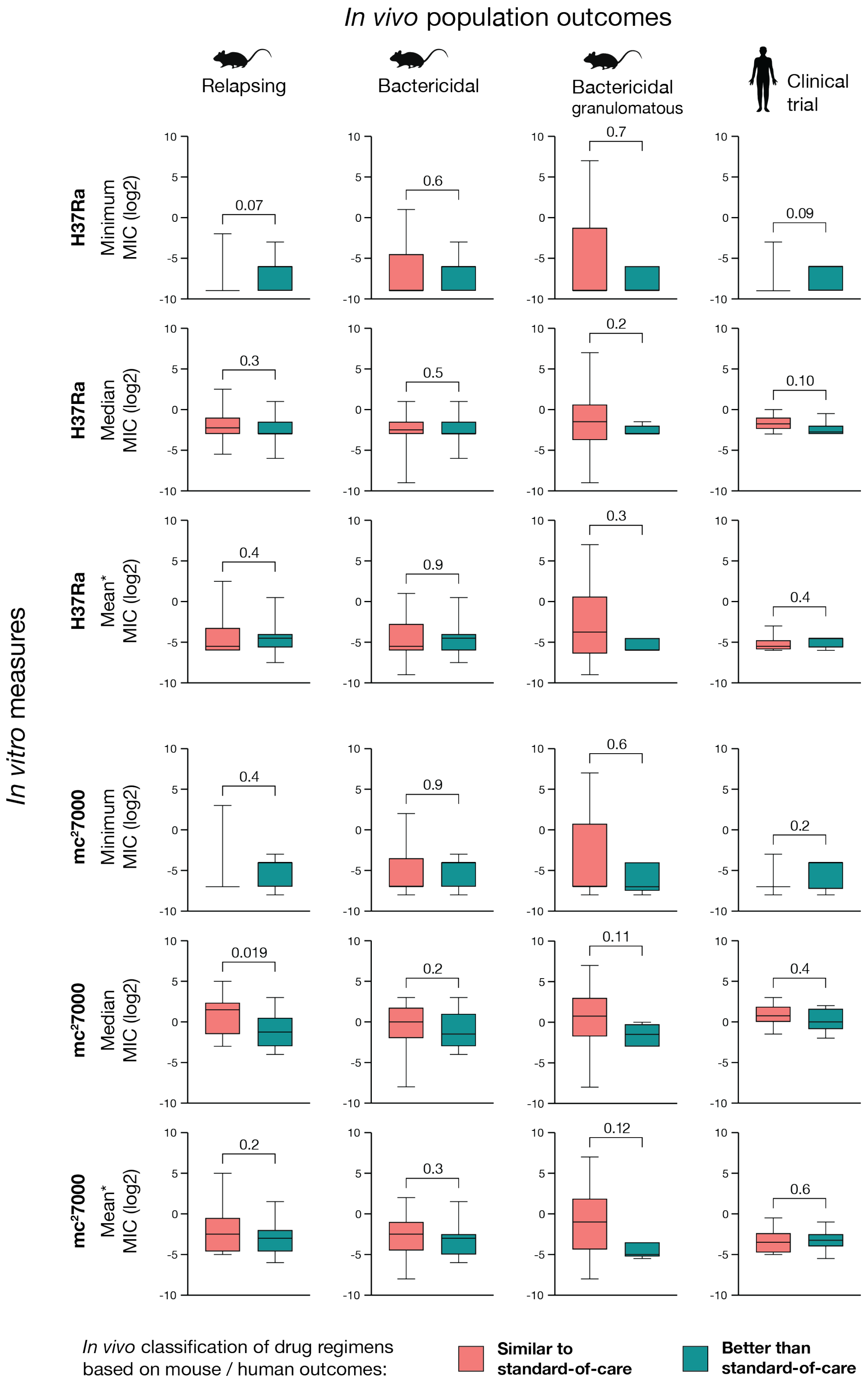

**Figure S4. Association of *M. tuberculosis* drug regimen MICs with *in vivo* outcomes.** *M. tuberculosis* drug regimens were previously classified as similar or better than standard of care (SOC) based on performance in relapsing mouse models (RMM), bactericidal mouse models (BMM) of common mouse strains, the granulomatous C3HeB/FeJ strain and clinical studies (*8*, *9*). The minimum MIC or median MIC of all drugs within each combination, or the mean MIC of the two most potent drugs (mean*), was assessed and compared between “similar to SOC” and “better than SOC” classifications using the Mann-Whitney U test (P indicated). Each dot represents a drug regimen. Boxplots show the median, interquartile range and total range of MIC values. (RMM: n = 46, BMM common strains: n = 48, BMM C3HeB/FeJ: n = 15, Clinical bactericidal activity: n = 14).

### Supplementary Figure 5

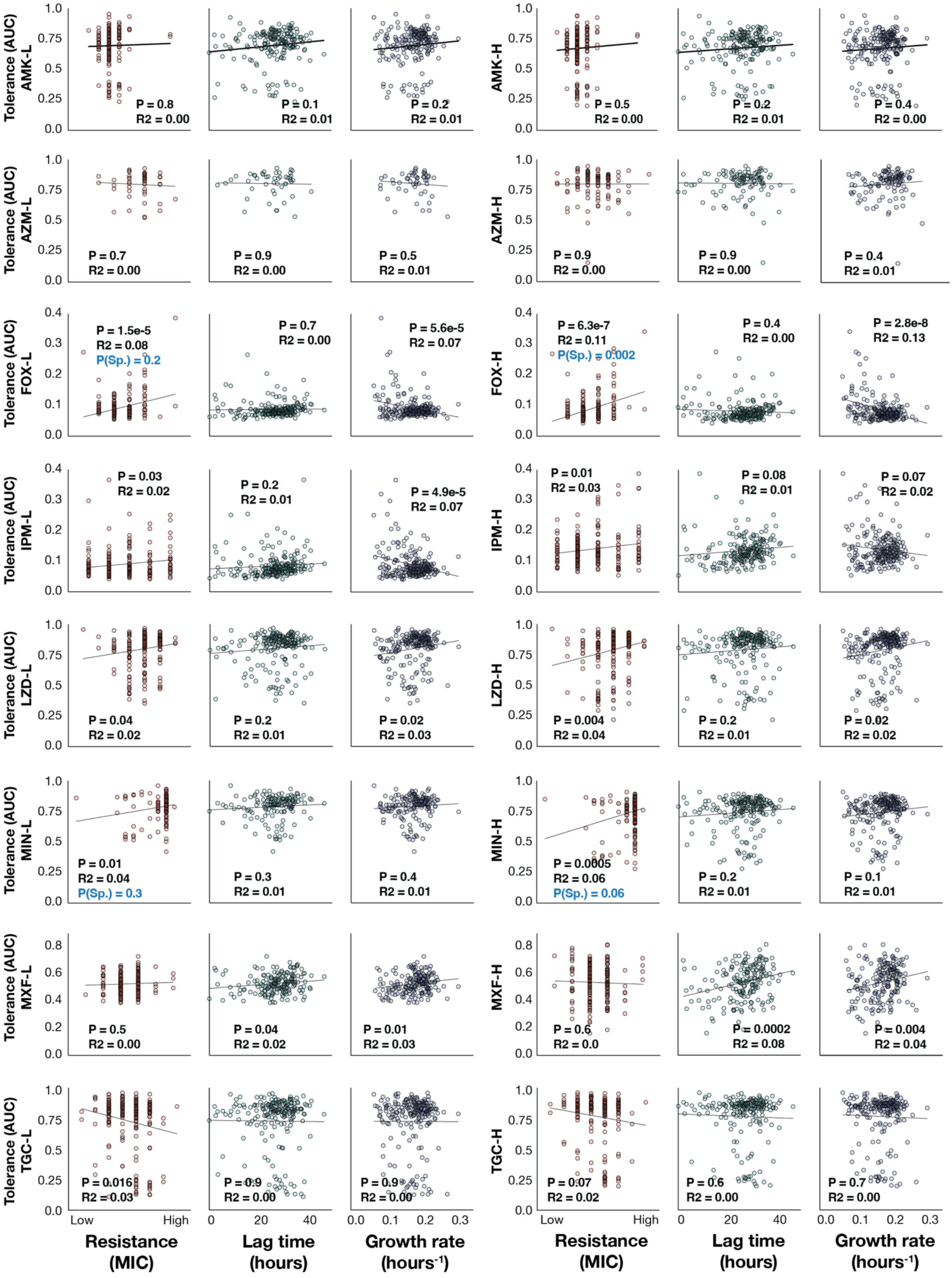

**Figure S5. Correlations of *M. abscessus* drug tolerance with bacterial replication and drug resistance.** Pearson correlation analyses (in blue Spearman correlation) between drug tolerance and bacterial growth rate, lag time, and the corresponding minimum inhibitory concentration (MIC). AUC indicates the area under the kill curve, AMK amikacin, AZM azithromycin, CLR clarithromycin, FOX cefoxitin, IPM imipenem, LZD linezolid, MIN minocycline, MXF moxifloxacin, TGC tigecycline, L refers to low drug concentration, and H to high drug concentration.

#### Supplementary Figure 6

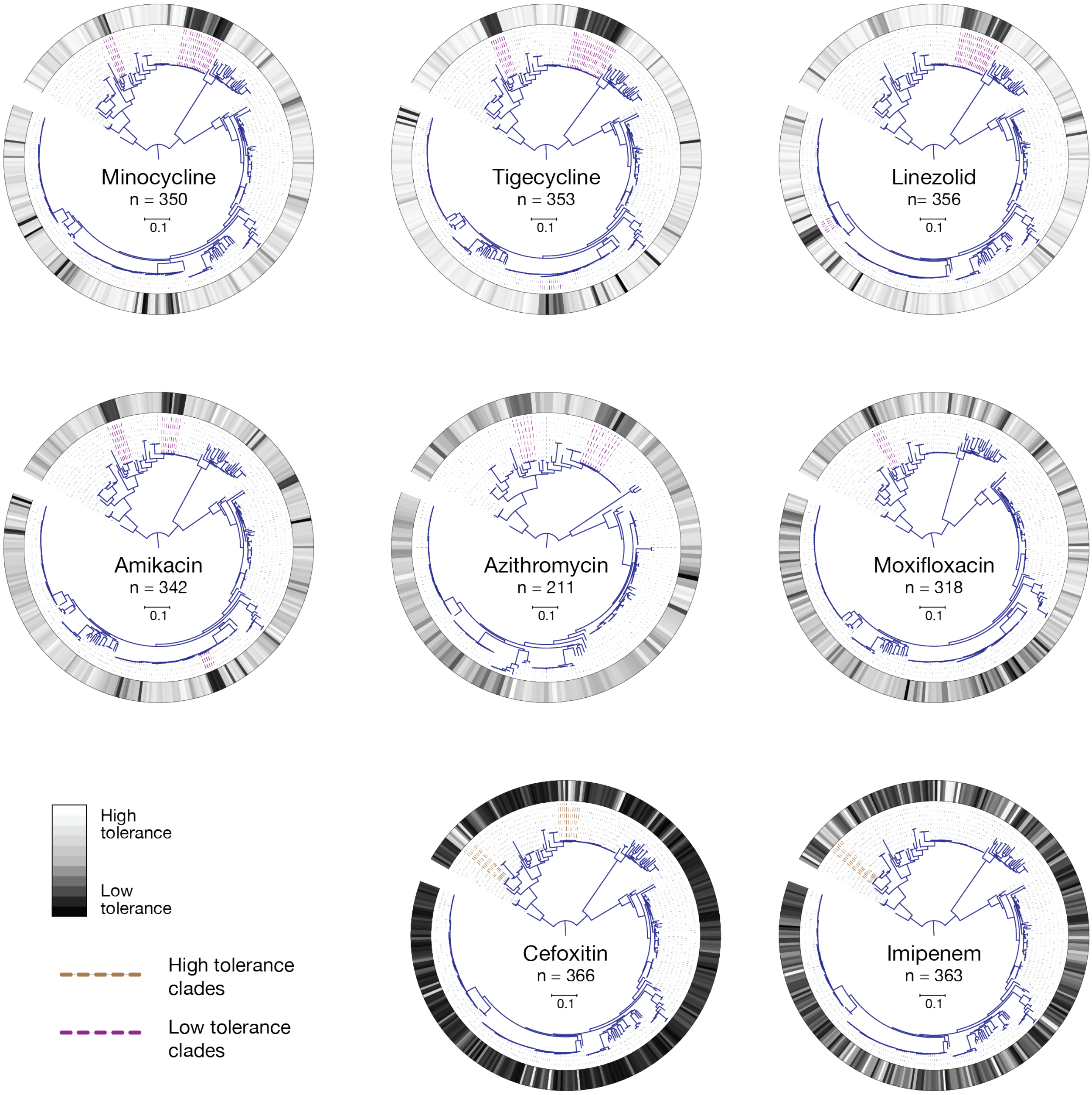


**Figure S6. Drug tolerance across the *M. abscessus* phylogeny.** *M. abscessus* phylogenetic trees (maximum-likelihood trees) aligned with drug tolerance heatmaps.

### Supplementary Figure 7

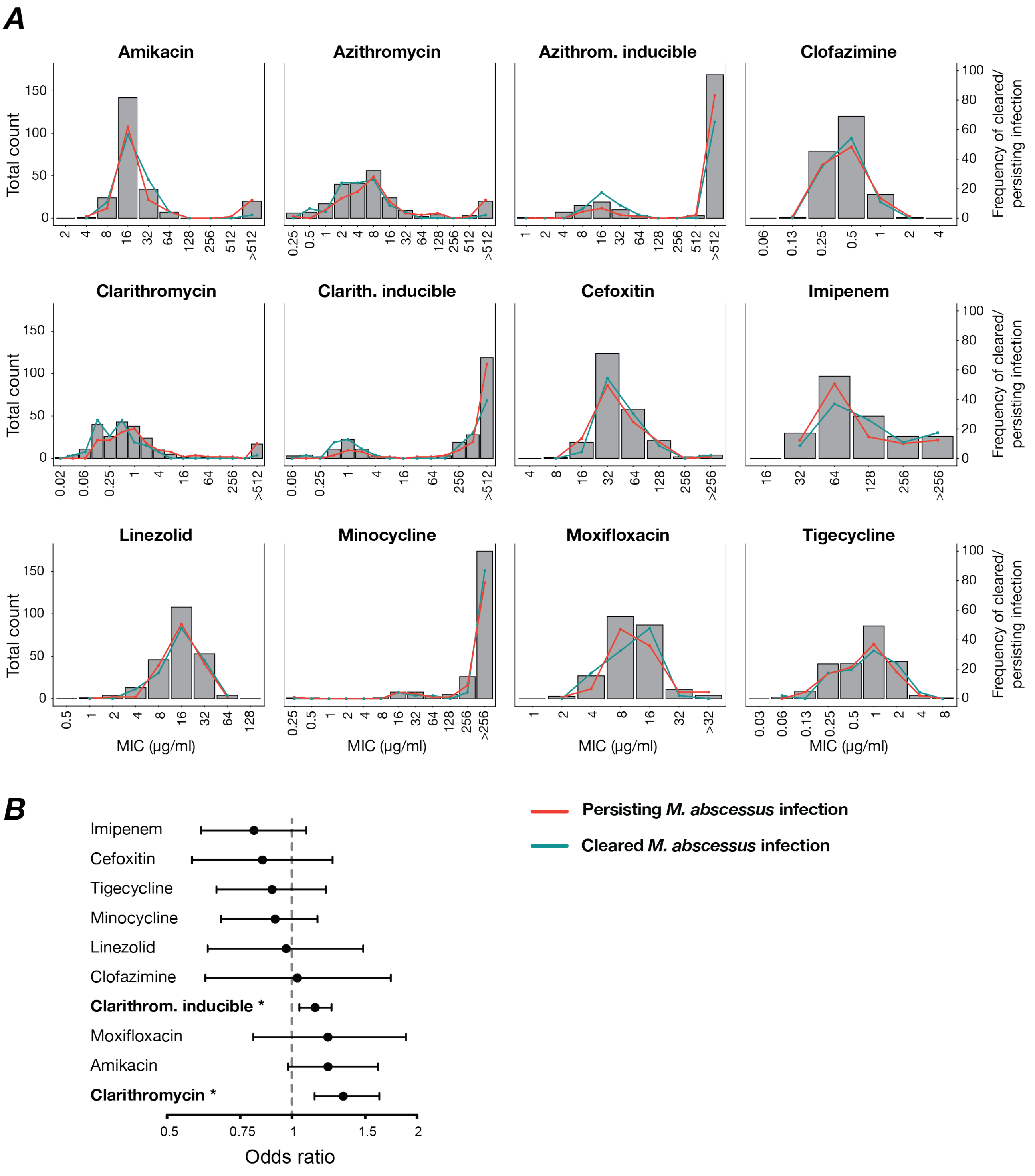


**Figure S7. Association of *M. abscessus* MICs with individual patient outcomes. (A)** MIC distributions of 229 clinical isolates and the frequency of isolates associated with cleared (n = 46) and persisting infection (n = 89). **(B)** Odds ratios of *M. abscessus* minimum inhibitory concentrations for predicting treatment failures (lack of culture conversion) in 135 patients. Odds ratios (95% CI) of individual MIC measures were assessed with logistic regression. * indicates P < 0.05.

#### Supplementary Figure 8

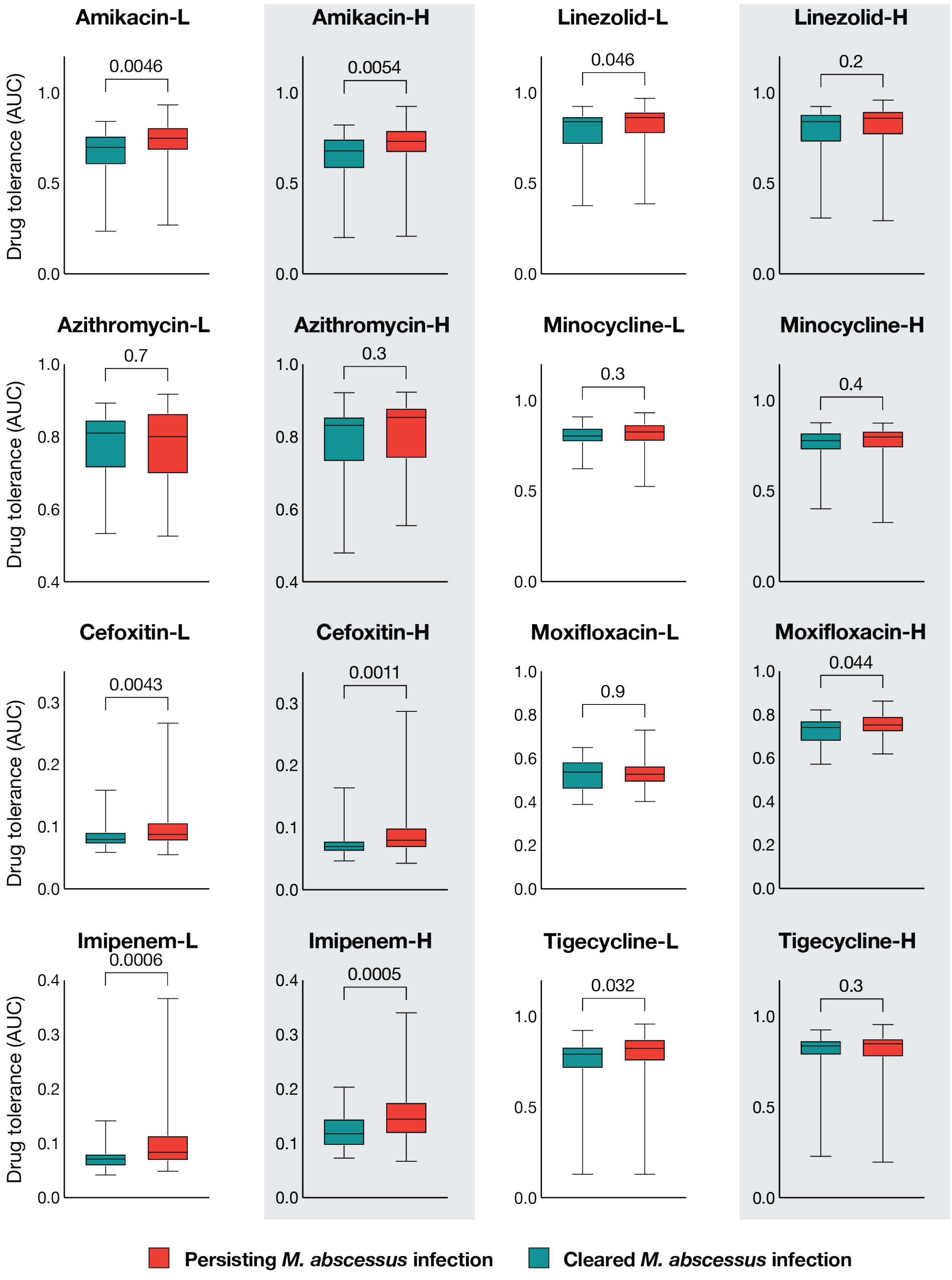

**Figure S8. Association of *M. abscessus* drug tolerance phenotypes with individual patient outcomes.** Comparison of drug tolerance between *M. abscessus* isolates from patients with poor versus favourable clinical outcomes, using the Mann-Whitney U test (P indicated). Boxplots show the median, interquartile range and total range of AUC values. L refers to low drug concentration, and H to high drug concentration.

#### Supplementary Figure 9

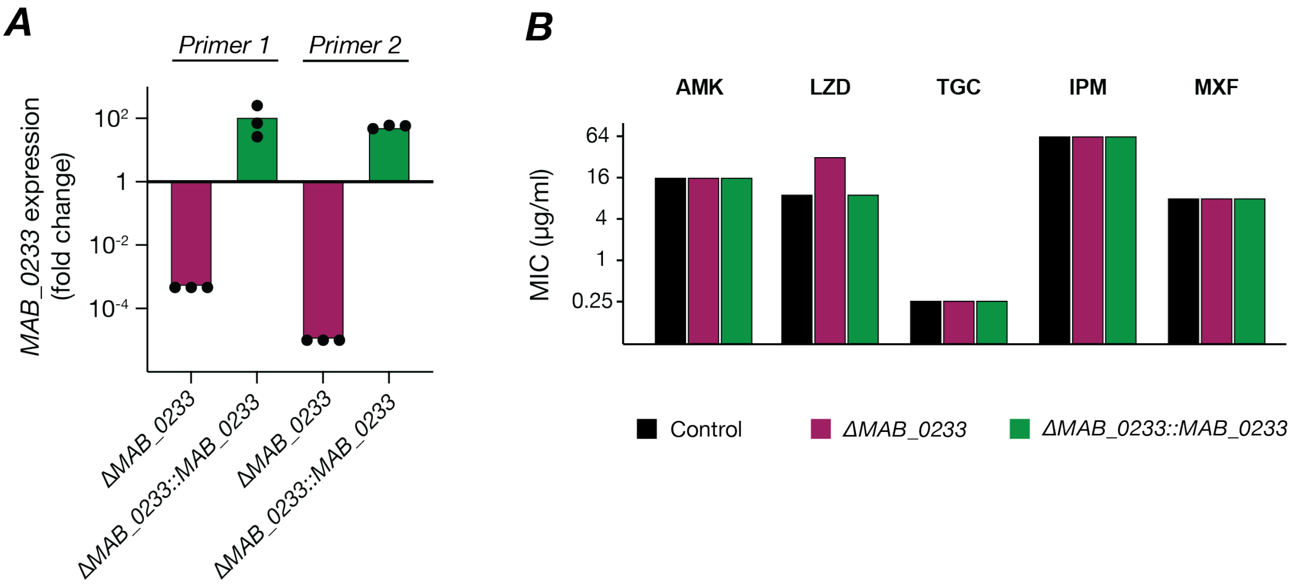


**Figure S9. Gene expression and minimum inhibitory concentrations of MAB_0233 mutants.** **(A)** Log-fold change of *MAB_0233* expression in the *MAB*_0233 knockout and complemented strain compared to the control *M. abscessus* strain during exponential growth. **(B)** Minimum inhibitory concentrations (MICs) of control, knockout and complemented strains. AMK indicates amikacin, IPM imipenem, LZD linezolid, MIN minocycline, TGC tigecycline.
